# A rectal cancer model establishes a platform to study individual responses to chemoradiation

**DOI:** 10.1101/640193

**Authors:** Karuna Ganesh, Chao Wu, Kevin P. O’Rourke, Mohammad Adileh, Bryan C. Szeglin, Isaac Wasserman, Michael R. Marco, Maha Shady, Youyun Zheng, Wouter R. Karthaus, Helen H. Won, Seo-Hyun Choi, Raphael A. Pelossof, Afsar Barlas, Emmanouil Pappou, Arthur Elghouayel, James S. Strong, Chin-Tung Chen, Jennifer W. Harris, Martin R. Weiser, Garrett M. Nash, Jose G. Guillem, Iris H. Wei, Andrea Cercek, Richard N. Kolesnick, Katia O. Manova-Todorova, Leonard B. Saltz, Ronald P. DeMatteo, Joan Massagué, Paul B. Romesser, Philip B. Paty, Rona D. Yaeger, Hans Clevers, Michael Berger, Jinru Shia, Scott W. Lowe, Lukas E. Dow, Julio Garcia-Aguilar, Charles L. Sawyers, J. Joshua Smith

**Affiliations:** Cancer Biology and Genetics Program, Sloan Kettering Institute, Memorial Sloan Kettering Cancer Center, New York, NY; Gastrointestinal Oncology Service, Department of Medicine, Memorial Sloan Kettering Cancer Center, New York, NY; Human Oncology and Pathogenesis Program, Memorial Sloan Kettering Cancer Center, New York, NY; Colorectal Service, Department of Surgery, Memorial Sloan Kettering Cancer Center, New York, NY; Department of Surgery, Hospital of the University of Pennsylvania, Philadelphia, PA; Gastrointestinal Pathology, Department of Pathology, Memorial Sloan Kettering Cancer Center, New York, NY; Department of Radiation Oncology, Memorial Sloan Kettering Cancer Center, New York, NY; Weill Cornell Medicine, Rockefeller University, Memorial Sloan Kettering Cancer Center Tri-Institutional MD-PhD Program, New York, NY; Marie-Josée and Henry R. Kravis Center for Molecular Oncology, Memorial Sloan Kettering Cancer Center, New York, NY; Molecular Pharmacology Program, Sloan Kettering Institute, Memorial Sloan Kettering Cancer Center, New York, NY; Molecular Cytology Core Facility, Memorial Sloan Kettering Cancer Center, New York, NY; Hubrecht Institute, Royal Netherlands Academy of Arts and Sciences, University of Medical Center Utrecht, The Netherlands; Sandra and Edward Meyer Cancer Center, Departments of Medicine and Biochemistry, Weill Cornell Medicine, Weill Cornell Graduate School of Medical Sciences, New York, NY

## Abstract

Rectal cancer (RC) is a challenging disease to treat that requires chemotherapy, radiation, and surgery to optimize outcomes for individual patients. No accurate model of RC exists to answer fundamental research questions relevant to individual patients. We established a biorepository of 32 patient-derived RC organoid cultures (tumoroids) from patients with primary, metastatic, or recurrent disease. RC tumoroids retained molecular features of the tumors from which they were derived, and their *ex vivo* responses to clinically relevant chemotherapy and radiation treatment correlate well with responses noted in individual patients’ tumors. Upon engraftment into murine rectal mucosa, human RC tumoroids gave rise to invasive rectal cancer followed by metastasis to lung and liver. Importantly, engrafted tumors closely reflected the heterogenous sensitivity to chemotherapy observed clinically. Thus, the biology and drug sensitivity of RC clinical isolates can be efficiently interrogated using an organoid-based, *in vitro* platform coupled with endoluminal propagation in animals.

## Introduction

Colon and rectal cancers are responsible for 50,000 deaths per year in the United States^1–3^. Rectal cancer (RC) is particularly challenging, as treatment after diagnosis is more complex compared to colon cancer^4^ due to tumor location in the pelvis and proximity to critical genitourinary organs. Rectal cancer invading the perirectal tissues or lymph nodes are treated with tri-modal therapy which consists of neoadjuvant chemoradiation (CRT), surgical resection, and chemotherapy^4^. Some RC patients respond completely to CRT alone and can avoid surgery entirely^5,6^, but others respond poorly and require radical surgery^7^. Prospective identification of patients who would achieve a complete response^7,8^ after neoadjuvant therapy alone would enable more tailored individual treatment regimens^9,10^ and thereby minimize potential harm from overtreatment. The heterogeneity in clinical response and the morbidity associated with radical surgery highlights the need for more sophisticated modeling to predict response to standard therapies.

Few cell lines have been derived from RCs^11–14^. Despite the fact that RC is treated differently from colon cancer by using tri-modal therapy in a neoadjuvant context, the preclinical development of treatments for rectal cancer has historically relied on colon cancer cell lines^15^, highlighting the need to develop RC-specific models. Furthermore, efforts to derive organoid “biobanks” have focused primarily on colon cancer specimens^16^, with a biorepository of rectal cancer tissue or organoids remaining an unmet need for the field.

An additional need for rectal cancer research is an anatomically accurate *in vivo* model using patient-derived RC organoids. The rectum has unique venous drainage via the iliac vessels that gives rise predominantly to lung metastases^17^ (69%), and less frequently liver metastases (20%), which are more commonly seen in colon cancers. Given recent success transplanting mouse colon cancer cells into the colon lumen by our group^18^ and others^19^, we set out to derive RC organoids (hereafter “tumoroids”) from resected or biopsied RCs and use them to establish *ex vivo* and *in vivo* RC model. We investigated the ability of such tumoroids to model the molecular and histologic features of human rectal cancers, as well as tumor initiation, invasion and metastasis. We also investigate whether our *ex vivo* and *in vivo* platforms could be used to correlate with treatment response in individual patients within a time frame that could potentially inform clinical treatment decisions.

## Results

### Human RC tumoroid derivation and characterization

With the goal of generating RC models that would reflect the biology of an individual patient’s tumor, we adapted existing strategies for 3D *ex vivo* tumor culture^20,21^ to generate RC tumoroids from pre-and post-treatment patient samples. Basic characteristics of all patients from whom we attempted tumoroid derivation are shown in **Extended Data Figure 1**. Overall, we established 32 RC tumoroids from 25 individual patients with an overall success rate of 76% (32/42). In addition to these RC tumoroids, we also generated 19 normal rectal organoids from normal adjacent tissue (see **Supplementary Table 1**). Of the ten failed tumoroid attempts, three were maintained for 6 weeks before senescing. As expected from previous work in colon cancer, RC tumoroids could be cultured in the absence of key growth factors^16^ (e.g., R-spondin, Wnt-3a, Noggin), whereas normal rectal organoids remained growth factor dependent^21^.

Since endoscopic biopsies are routinely performed in the outpatient setting in the treatment of patients with rectal cancer, we asked whether we could derive RC tumoroids from minute amounts of material obtained in the clinic. Of note, 17 of the RC tumoroids were established using tissue obtained with biopsy forceps routinely used in clinical care (**Fig. 1a**). These data suggest it is possible to use serial biopsies obtained as part of standard care in the pre-treatment or post-therapy settings to generate tumoroid models to assess patient-specific mediators of response and resistance.

**Fig. 1.**
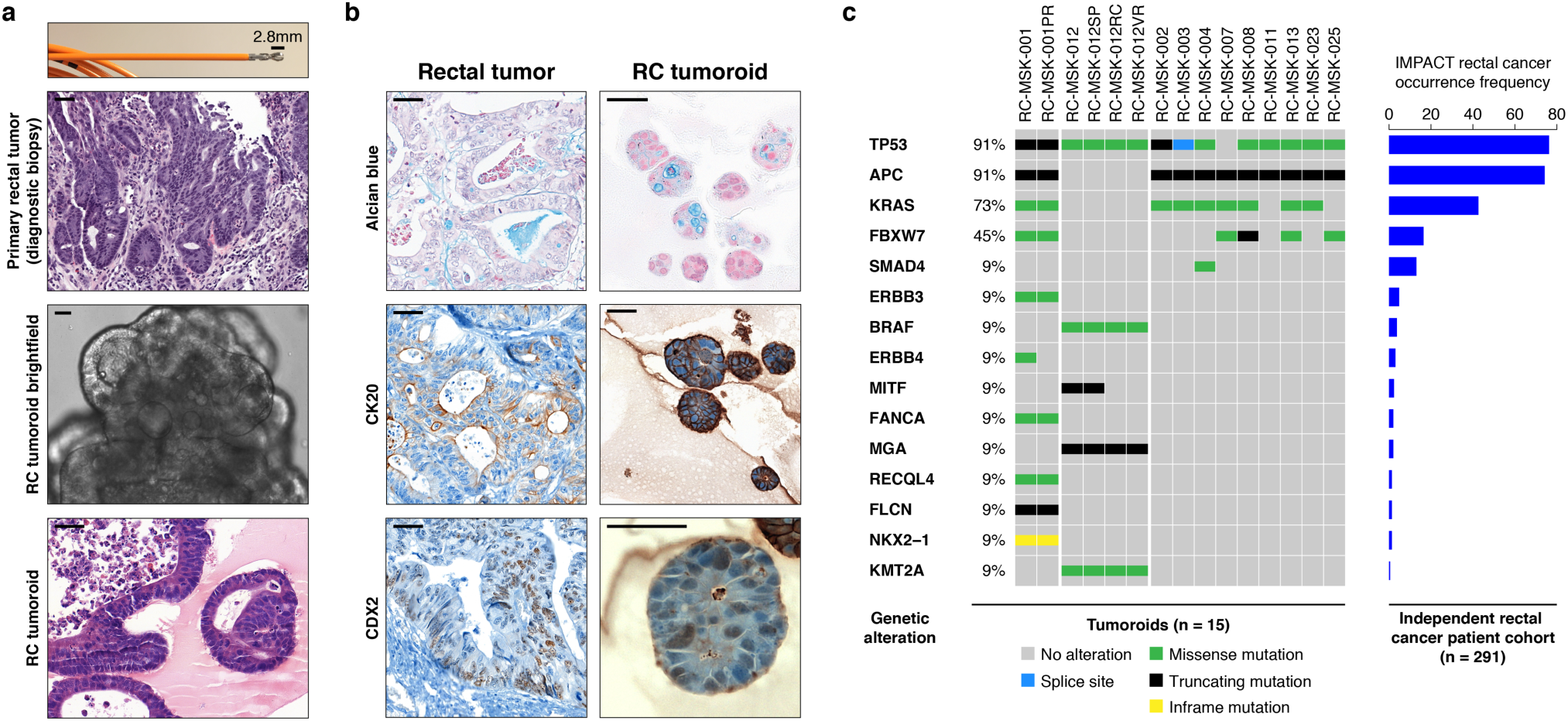
Preservation of rectal cancer histopathology and mutational fingerprint in tumoroids. **a**, Shown at the top is the 2.8 mm cold biopsy forceps (Boston Scientific Corp.™) used for sampling tumor from some of the rectal cancers (RCs) used to derive tumoroids. Also shown is the first primary tumor sampled with this biopsy forceps stained with hematoxylin and eosin (H&E, second panel). The corresponding derived tumoroid, RC-MSK-008, in 3D culture is displayed by brightfield microscopy and H&E (lower 2 panels). Scale bars, 50 μm. **b**, Histopathologic staining of enterocyte markers (Alcian blue, CK20, and CDX2) of a primary resected rectal tumor (leftmost panels) and the corresponding tumoroid, RC-MSK-001, in 3D culture (right panels). Scale bars, 50 μm. **c**, The mutation landscape of 15 of the RC tumoroids is displayed (left panel) compared to an independent set of 291 RCs (right panel), both detected by MSK-IMPACT. The frequency of alterations in the RC tumoroids is noted with the type of genetic alteration (noted by color code). The figure shows the top 15 mutated genes in the 291 patient cohort are observed in the derived tumoroids.

Despite concern that tumor stem cells exposed to prior systemic chemotherapy or radiation would not likely yield viable tumoroids^16^, we were able to generate RC tumoroids and normal adjacent rectal organoids from patients who had undergone prior chemotherapy and/or radiotherapy, as well as those who had no prior therapeutic intervention before surgical resection. Of the 32 tumoroids, 9 were derived from treatment-naïve patients, 17 from patients undergoing first or second line therapy (**Supplementary Table 1**). The remaining 6 were derived from patients with sites of disease recurrence: 4 perineal local recurrences and 2 metastases (a paired splenic metastasis and a peritoneal metastasis). Tumoroids were derived from all sites of the rectum (i.e., distal, mid and upper), from *RAS* mutant (64%) and wild type tumors, and from patients with metastatic disease (48%) and non-metastatic disease (**Extended Data Fig. 1**). In summary, we sampled a diverse set of RC patients with different stages of disease, on or off therapy, *RAS* mutant and wild type status, and did not find any clinical variables that predicted success or failure of organoid establishment.

### Histopathologic and molecular characteristics are preserved in rectal cancer tumoroids

As with cancer organoids derived from other tissues, RC tumoroids retained histopathologic features of the primary tumors, such as preservation of the glandular structure and lumen (**Fig. 1a, Extended Data Fig. 2**). Further, RC tumoroids retained Alcian blue-positive and MUC-2-positive goblet cells, CK20 and CDX2-positive enterocytes, and robust expression of E-cadherin (epithelial marker), mirroring the tumors from which they were derived (**Fig. 1b, Extended Data Fig. 3**). The tumoroids demonstrate variable CDX2 staining, as seen in the primary tumor. These data indicate that features found in the primary tumor are retained in the tumoroid from which it was derived and reflect individual patient tumors.

To determine whether the RC tumoroids also reflect the mutational fingerprint in the corresponding primary tumors—as well as the larger landscape of molecular alterations in RC—we performed massively parallel sequencing using the custom, FDA-cleared tumor profiling panel MSK Integrated Mutation Profiling of Actionable Cancer Targets (MSK-IMPACT)^22^. We sequenced genomic DNA isolated from 15 of the tumoroids and the corresponding patient tumors. Tumoroid mutations are summarized in **Extended Data Figure 4a**. The degree of conservation of mutations between examples of tumoroids and their primary tumors is shown in **Extended Data Figure 4b and c** and shown for all tumoroids in **Extended Data Figure 5**. Among mutations likely to be oncogenic, concordance was 87% (range 0.49-1.00; **Extended Data Figure 5b**), which is comparable to an 88% concordance (range 0.62-1.00) previously reported for colon organoids^16^ and organoids from colon metastases^23^ (no rectal cancers). Of the clonal mutations present in the matched tumor samples, 73% were also present as clonal in tumoroid samples. We then compared these tumoroids to a clinical cohort of 291 rectal cancers resected at Memorial Hospital that had been sequenced by MSK-IMPACT. The top mutations in the 291 rectal cancer patients were also noted in the RC tumoroids, with the most common alterations in the genes *APC*, *TP53, KRAS*, and *FBXW7* (**Fig. 1c**).

### RC tumoroids reveal a diversity of responses to chemotherapy that correlate with clinical response

We next examined our RC tumoroids as a platform for preclinical studies using the established RC tri-modal treatment approaches. 5-fluorouracil (5-FU) is the backbone of RC chemotherapy as part of systemic therapy or in combination with radiation^24,25^. Treatment of six different RC tumoroids with 5-FU yielded significant differences in cell death (**Fig. 2a**; Kruskal-Wallis test for nonparametric data, P < 0.02) and heterogeneous IC_50_ values (range: 34-800 μM, **Fig. 2b**, right axis). Within the therapeutic dosage range noted clinically^26^ (19-23 μM, shown as grey box in **Figure 2a**), we observed significant differences in cell death among the tumoroids. Thus, the RC tumoroids are heterogeneous in their response to physiologic 5-FU doses, suggesting that they could be used as a tool to predict patient response to 5-FU treatment.

**Fig. 2.**
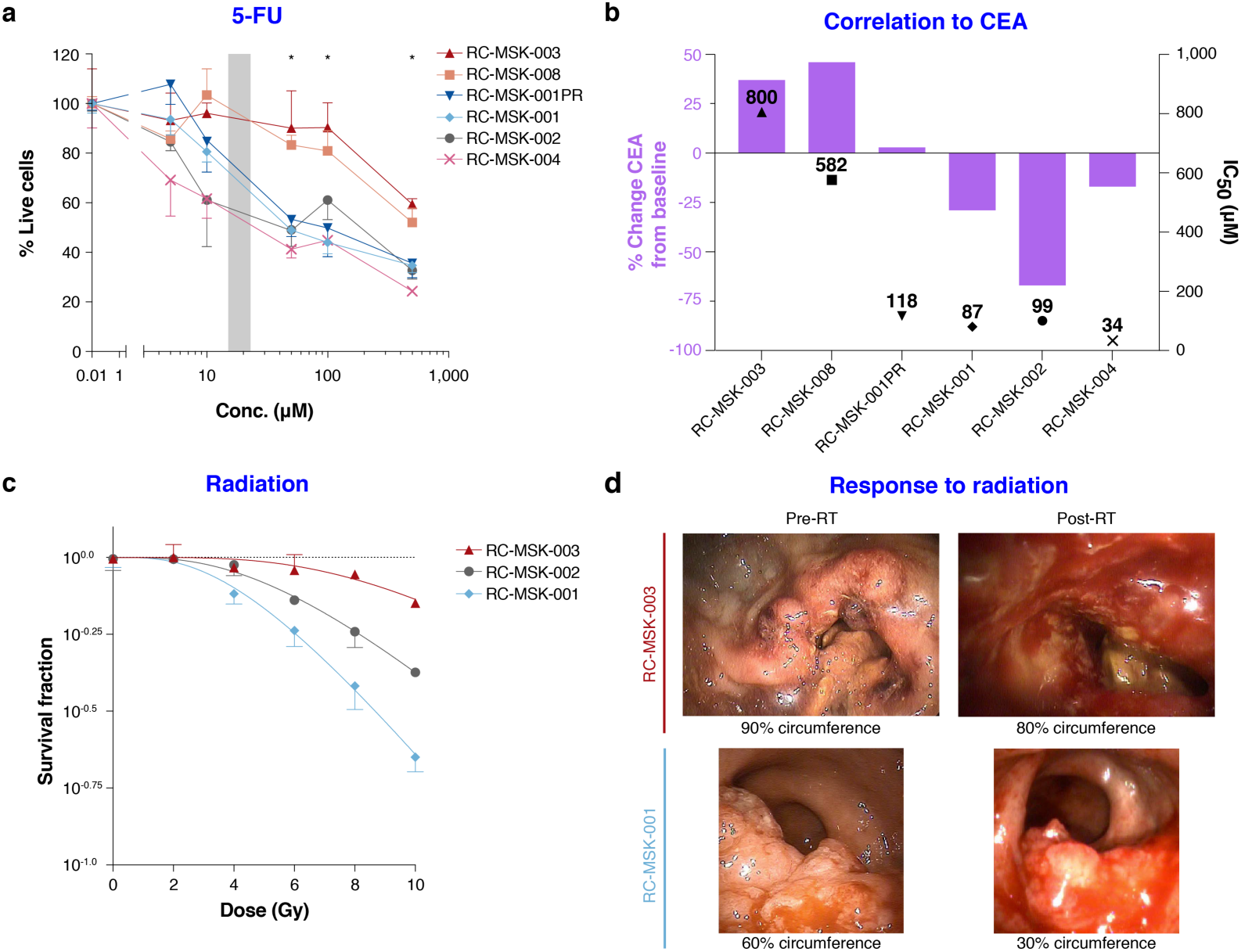
Clinically relevant responses to chemotherapy and radiation in rectal cancer tumoroids ex vivo. **a**, Sensitivity of RC tumoroids to 5-fluorouracil (5-FU) doses. The percent live cells for each tumoroid is displayed (data in quadruplicate, error bars represent standard error of the mean [SEM]). The grey box represents the therapeutic dosage range for 5-FU: 19-23 μM^26^. Statistical differences were analyzed by the Kruskal-Wallis test for each drug concentration. P-values are annotated as: * < 0.02. We note the details of the Kruskal-Wallis test statistic [H], degrees of freedom [DF] in the methods and the specific P-values here: 0.01 μM: P = 0.971; 5 μM: P = 0.150; 10 μM: P = 0.067; 50 μM: P = 0.005; 100 μM: P = 0.014; 500 μM: P = 0.0059. **b**, IC_50_ values for RC tumoroids (right axis) juxtaposed with a bar graph displayed as a modified waterfall plot (left axis) showing treatment response of the corresponding patients as percent change in carcinoembryonic antigen (CEA, tumor marker, left axis) from baseline. **c**, Sensitivity of three RC tumoroids to ionizing radiation is shown as the surviving fraction per dose in Gy across a range of radiation doses. D_0_ and D_q_ for each are as follows (RC-MSK-003: D_0_ = 11.00 and D_q_ = 6.31; RC-MSK-002: D_0_ = 6.70 and D_q_ = 4.93; RC-MSK-001: D_0_ = 5.33 D_q_ = 2.72) (data in triplicate; error bars = SEM). **d**, Clinical responses for patients from whom RC-MSK-003 and RC-MSK-001 were derived. Displayed are endoscopic photos taken in the clinical setting pre- and post-radiation therapy (RT). RC-MSK-003 primary tumor (top panels) showed minimal change (bleeding and minor reduction (10%) in the circumferential measurement of the tumor). RC-MSK-001 primary tumor (bottom panels) showed a 50% reduction in tumor size as noted by circumferential endoscopic measurement. The RC-MSK-002 patient was not treated with radiation and so is not shown.

We then asked whether *ex vivo* 5-FU sensitivity correlated with a parameter of clinical utility in the corresponding patients as assessed by carcinoembryonic antigen (CEA), which is a marker of active disease or recurrent disease^27,28^. The change in CEA levels from baseline for the RC-MSK-003 and RC-MSK-008 patients with higher IC_50_ values was larger (+37% and +48%) than it was for the RC-MSK-001PR and RC-MSK-001 (+2.9% and −29%) and the RC-MSK-002 and RC-MSK-004 patients (−67% and −17%) (**Fig.2b**, Pearson’s Correlation: r = 0.801, P = 0.055). These data also suggest that 5-FU sensitivity determined *ex vivo* could have utility in predicting clinical sensitivity.

### Ex vivo irradiation response correlates with clinical response

RC-MSK-001 showed significant response to radiotherapy (RT, D_0_ = 5.33), while RC-MSK-002 (D_0_ = 6.70) and RC-MSK-003 (D_0_ = 11.0) were resistant to radiation *ex vivo* (**Fig. 2c**). *Ex vivo* RT sensitivity of the RC-MSK-001 and RC-MSK-003 tumoroids correlated with clinical RT sensitivity of the tumors from which they were derived as assessed by changes in luminal circumference of the tumor by endoscopy before and after radiation (**Fig. 2d**: RC-MSK-001, 60% of circumference to 30%; RC-MSK-003, 90% of circumference to 80%). The patient from which RC-MSK-002 was derived was not treated with neoadjuvant radiation. Taken together, these data demonstrate that RC tumoroids display varying sensitivity to ionizing radiation, indicating that this platform could be used to evaluate patient response before initiating this treatment modality and potentially sparing resistant patients the toxic side effects of radiation therapy^29^.

### Proof-of-principle targeted therapy studies in RC tumoroids

In colorectal cancers, the presence of *KRAS* mutation predicts resistance to EGFR-targeted therapy; therefore, we asked whether RC tumoroids would recapitulate *KRAS* mutation-mediated resistance to the anti-EGFR monoclonal antibody cetuximab^30^. Consistent with clinical data, *KRAS*^mutant^ tumoroids were resistant to cetuximab, whereas *KRAS*^wild type^ tumoroids were sensitive (**Extended Data Fig. 6**). These data provide preliminary evidence that a cell-autonomous response in RC tumoroids reflects a clinically important response to a targeted therapy that could readily be assayed *ex vivo* to functionally determine individual patient responses.

### Establishing an endoluminal model using human rectal cancers

We next sought to establish an orthotopic, patient-derived xenograft model of RC that could reflect the initiation, invasion, and metastatic potential of tumors that develop in the unique anatomy of the rectum. **Fig. 3a** shows the methodology^18^ and documents engrafted human rectal cancer in a NOD scid gamma (NSG) mouse rectum after injecting 200,000 cells from tumoroids (**Supplementary Movie 1**, online). Successful transplantation of human tumoroids was confirmed by human EpCAM staining of an endoscopic biopsy specimen taken from the live mouse compared with negative staining in the NSG proximal mouse colon as a control (**Extended Data Fig. 7**). **Figure 3a** also highlights serial assessment of engrafted rectal tumors at 8 and 12 weeks by endoscopy. We noted progression to intramucosal adenocarcinoma at 16 weeks post-transplantation in our first experiments (**Fig. 3b** top panel). The endoluminal rectal cancer model was reproducible in both male and female mice using cells from six different RC tumoroid lines. Monthly endoscopy assessed all lesions (**Extended Data Fig. 8a and b**) and gross tumor formation ranged from 20%-100% per experiment. Even though we did not observe gross tumors in all mice, 100% of the mice transplanted showed engraftment of cells positive for human EpCAM (**Extended Data Fig. 8c and d**). We also demonstrated that tumoroids can be genetically modified prior to transplantation, which permits tracing or mechanistic studies, as demonstrated by GFP-labeling of tumoroids *in vitro* prior to endoluminal transplantation (**Extended Data Fig. 8e and f)**.

**Fig. 3.**
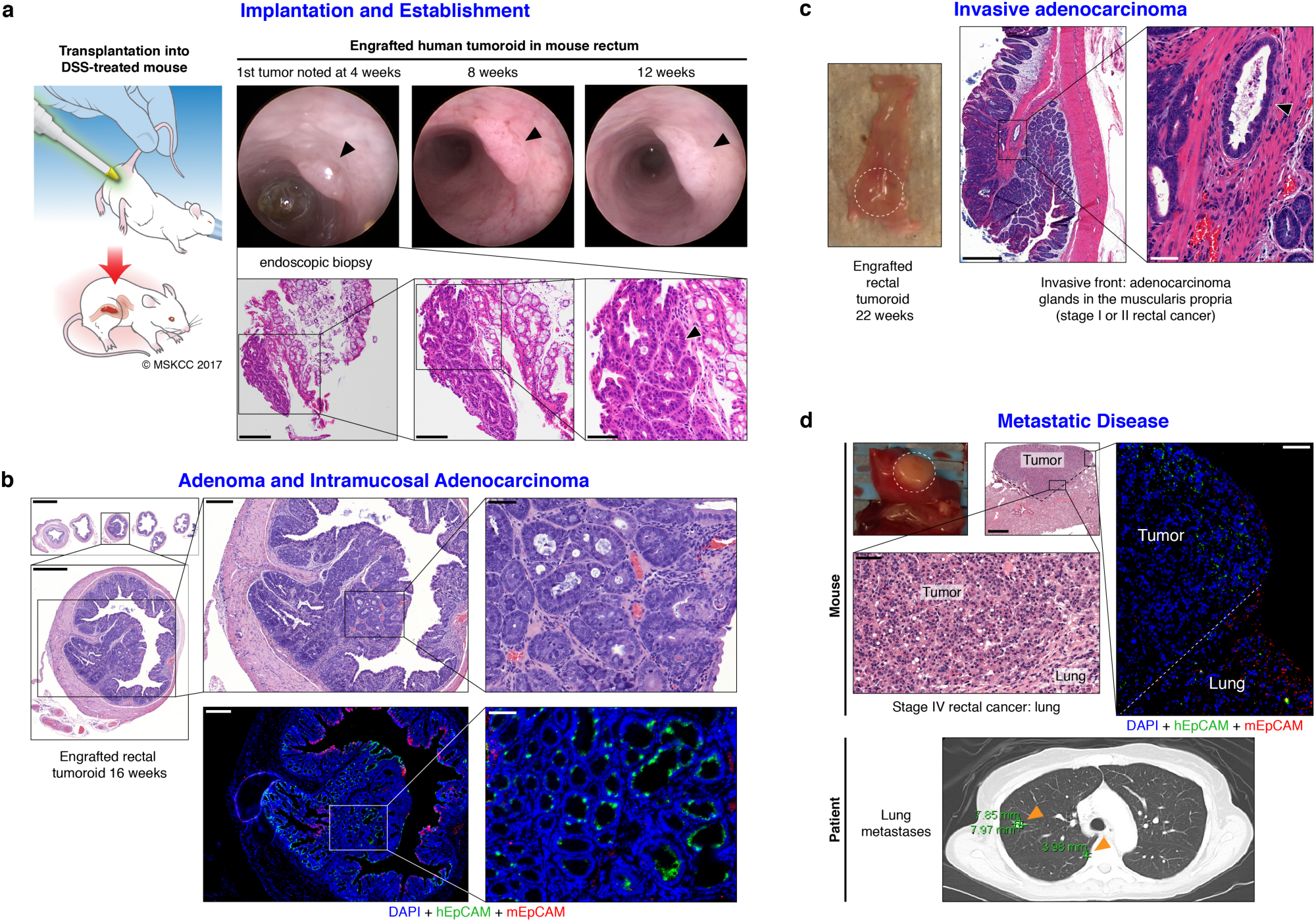
Establishment of an endoluminal rectal cancer assay in mice. **a**, Cartoon of the implantation process (T. Helms) and an endoscopic view of the first rectal tumoroid (RC-MSK-001) implanted in a NOD scid gamma (NSG) mouse at 4, 8, and 12 weeks (top panels). H&E of the engrafted tumoroid biopsied at 4 weeks (December 2015) via endoscopic channel (bottom panels) demonstrates high nuclear/cytoplasmic ratio, poor differentiation, and neoplastic glands (arrowhead). Adjacent normal mouse colonic epithelium and stroma is seen. Scale bars from low to high magnification are as follows: 200 μm, 100 μm, 50 μm. **b**, H&E, axial view of the rectal tumor within the NSG mouse rectum with adjacent serial sections (upper panels). Inset demonstrates intramucosal adenocarcinoma with atypical neoplastic cells, poorly differentiated tumor cells, and high nuclear to cytoplasmic ratio. Immunofluorescence (IF, lower panels): DAPI (blue), human (h) EpCAM (green), mouse (m) EpCAM (red). Scale bars from low to high magnification: 2,000 μm, 500 μm, 200 μm, 50 μm. **c**, NSG mouse rectum (gross rectum with tumor, white dashed circle) and H&E shows evidence of an engrafted human rectal tumor with invasion into the muscularis (inset, arrowhead). Scale bars: low magnification, 500 μm; high magnification inset, 50 μm. **d**, Gross lung metastasis (white dashed circle) from an engrafted human RC tumoroid and corresponding H&E are shown. IF serial section demonstrates engraftment of the human tumoroids: DAPI (blue), hEpCAM (green), mEpCAM (red). Lower panel shows axial thoracic CT imaging of the patient from which the tumoroid was derived, demonstrating lung metastases (orange arrows indicate two of multiple lung metastases). Scale bars: low magnification, 500 μm; H&E and IF high magnification insets, 50 μm.

We then interrogated our model for to the development of invasive cancer and metastatic disease. At 22 weeks and 30 weeks post-transplantation, we noted progression to invasive adenocarcinoma that parallels the features of stage I or II human rectal adenocarcinomas (**Fig. 3c** and **Extended Data Fig. 9**). Importantly, we noted evidence of metastases in the lung (**Fig. 3d**) and liver (**Extended Data Fig. 10a**) that demonstrated features of poorly differentiated carcinomas infiltrating normal parenchyma, as confirmed by independent pathological review. Notably, the metastases derived from the endoluminal rectal tumors corresponded to sites of metastases seen in the individual patients from which the tumoroids were derived, namely lung metastases in the case of RC-MSK-001 and liver for RC-MSK-002 (**Fig. 3d** lower panel, **Extended Data Fig. 10b**, respectively). The metastases were further confirmed with human-specific PCR analyses (**Extended Data Fig. 10c**). These data indicate the feasibility of a patient-derived RC endoluminal model and its patterns of clinically relevant metastasis.

### The endoluminal model as a platform to investigate chemosensitivity

We hypothesized that we could mimic chemotherapy response in our *in vivo* RC model as a parallel, pre-clinical assay to our *ex vivo* work. To establish safe use of 5-FU *in vivo*, we first conducted experiments in a subcutaneous injection model and treated the mice systemically with 50 mg/kg/week of intraperitoneal injection 5-FU. The tumoroids successfully engrafted, the mice tolerated treatment well, and we noted a level of response in two separate tumoroid lines (**Extended Data Fig. 11**).

After showing safe use of 5-FU *in vivo*, we then tested whether we could demonstrate chemosensitivity in our endoluminal RC model. We implanted mice with tumoroids from two lines: RC-MSK-008, which was 5-FU–insensitive both clinically and *ex vivo*, and RC-MSK-004, which was 5-FU–sensitive in both circumstances (**Fig. 2a, b** and **Fig. 4a,c** bottom panels). At 12 weeks, 70% of the mice had tumors detectable by endoscopy. All mice were then randomized to 5-FU or vehicle treatment and the tumors were followed endoscopically every 3-4 weeks. Mice engrafted with RC-MSK-008 tumoroids demonstrated a lack of response to treatment with 5-FU (**Fig. 4a and b**). In contrast, mice engrafted with RC-MSK-002 tumoroids demonstrated significant reduction in tumor growth post-5-FU treatment (**Fig. 4c and d**). Of note, these observations were consistent with the individual patient’s clinical responsiveness to 5-FU-based systemic chemotherapy. The tumoroid-derived endoluminal rectal xenografts were confirmed histologically to represent the glandular morphology of the primary tumors from which they were derived (**Extended Data Fig. 12**). These experiments establish the feasibility of an *in vivo* chemosensitivity assay using the endoluminal model, which could be used to corroborate *ex vivo* drug resistance and, more importantly, provide a relevant platform for testing new agents in the context of resistant or sensitive tumors in a pre-clinical setting.

**Fig. 4.**
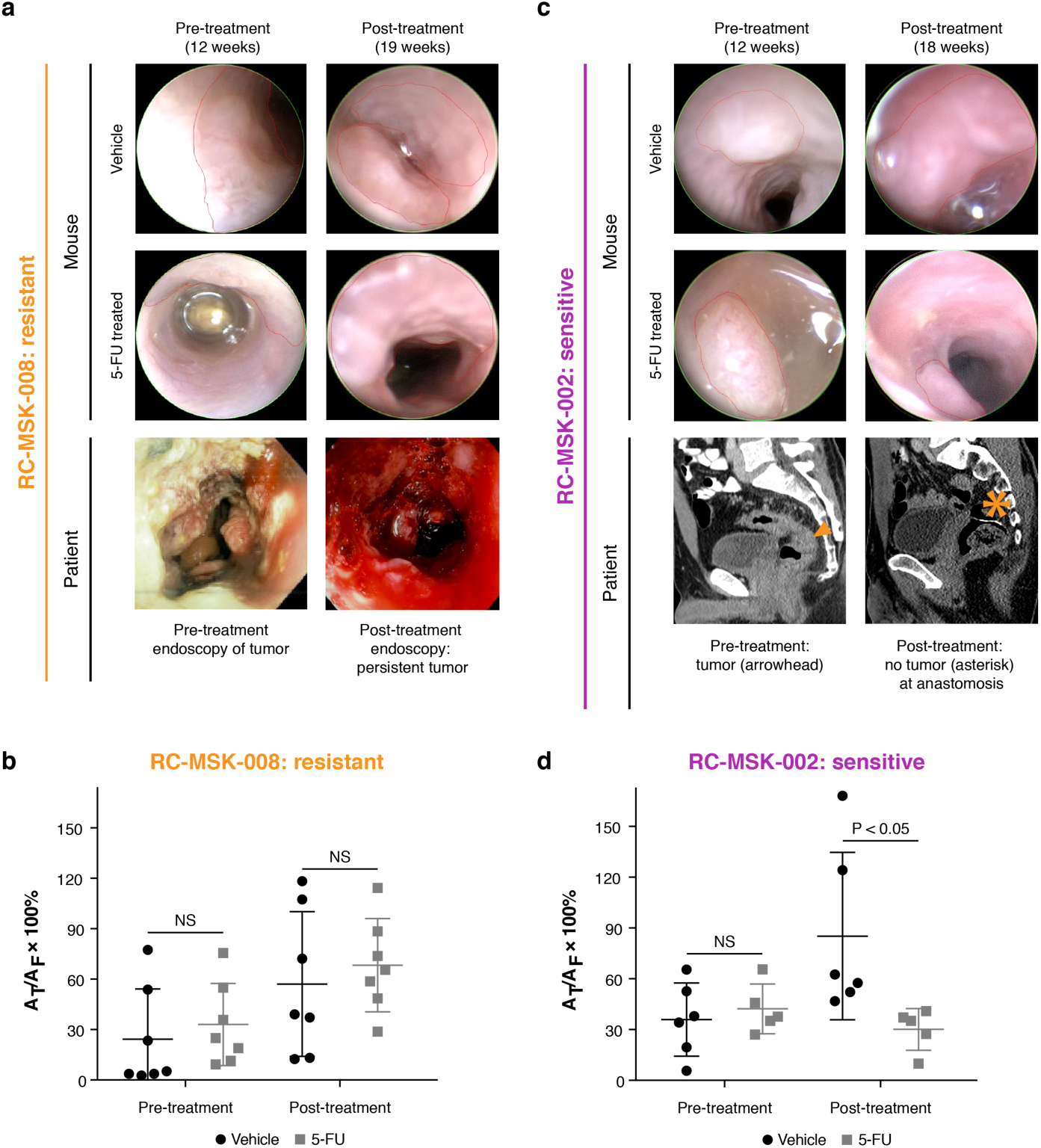
The endoluminal rectal cancer model can be used to reflect patient-specific chemoresistance and chemosensitivity. **a**, Upper two rows, RC-MSK-008 implanted in mice is shown pre- and post-treatment with vehicle or 5-FU. Tumor is outlined in red. Bottom row, the corresponding human rectal tumor viewed by endoscopy in the clinic at diagnosis and post-5-FU based treatment. **b**, Quantification of RC-MSK-008 tumor area measured endoscopically in the mice pre- and post-treatment (n = 7 per condition). Tumor growth is presented as area of tumor (At) as a percentage of area of field of view (Af), and was not significant (NS, P>0.05). This observation reflected the clinical response as noted in panel a (bottom row). Error bars: larger, middle lines represent the sample means; smaller, end lines represent sample standard deviations. **c**, Upper two rows, RC-MSK-002 implanted in mice is shown pre- and post-treatment with vehicle or 5-FU. Bottom row, sagittal CT imaging of the corresponding patient tumor pre-treatment (orange arrowhead: tumor) and after resection and 5-FU based systemic therapy (orange asterisk: anastomosis) with no evidence of recurrence. **d**, Quantification of RC-MSK-002 tumor area as in b. These data reflect the clinical scenario relative to the rectal tumor for this patient as noted in panel c (bottom panel). Tumors in b and d were confirmed histologically (**Extended Data Figure 12**). Error bars: as noted above for panels a,b.

## Discussion

We have established a biorepository of RC models representing the spectrum of genetic diversity and treatment response seen in RC. This unique resource represents a significant expansion of the reagents for the study of RC and will enable mechanistic and translational studies to be performed on bona fide RCs, rather than surrogate colon cancer cell lines. Importantly, RC tumoroids can be routinely and successfully established from small amounts of starting material using a 2.8 mm biopsy forceps, demonstrating that only a small amount of tissue is required to derive a viable tumoroid line and that samples can be obtained in the outpatient clinic. We have demonstrated that RC tumoroids from patients successfully engraft into the rectal mucosa of mice and that these orthotopic, endoluminal rectal xenograft mouse models recapitulate patient-specific treatment response *in vivo*. Notably, we contribute new methodology and modeling with regard to *ex vivo* radiation response and patient-specific *in vivo* chemotherapy response in our endoluminal model. To our knowledge, the correlation demonstrated here with radiation in RC tumoroids is the first demonstration of this *ex vivo* technology as it relates to clinical response in a disease where radiation is standard of care. Further, our data suggest that the *ex vivo* and *in vivo* models we have derived are a representative analog of individual RC patient response, complementing what has been recently described for other metastatic adenocarcinomas^31^.

Despite concerns that organoid technology would be unsuccessful with heavily pre-treated tumor specimens^16^, our findings are especially important as our work shows that this is possible with a success rate of ~75%. We have demonstrated that the basic histopathology and enterocyte differentiation markers are conserved between RC tumoroids and primary tumors. The mutational profiles between tumor and derived tumoroid were conserved and we noted similarity between our derived RC tumoroids and an independent MSK-Rectal Cancer Cohort (**Fig. 1**).

Recapitulation of patient-specific and clinically relevant responses to chemotherapy, irradiation, and targeted therapy may open doors to determine new and more precise treatments for RC patients. Furthermore, the derivation of RC tumoroids and normal rectal organoids from biopsies is critically important, as this means models can be derived at diagnosis or at the time of recurrence from minute amounts of biopsy material. *Ex vivo* and *in vivo* sensitivity to standard and experimental therapeutics (e.g. trials), as well as irradiation, could be fully assayed in 6-12 weeks after establishment (**Figs. 2 and 3**). If the individual patient fails standard treatment or experiences progressive disease, these assays may provide rationale to attempt new therapies found to be effective both *ex vivo* and *in vivo* using the patient’s own tumor tissues as the embedded informant. These data could be gathered within the period it takes a modern-day RC patient to undergo total neoadjuvant therapy (6 months)^32^. Future work will leverage our model to prospectively evaluate the concordance between patient and tumoroid response to standard therapeutics, including both chemotherapy and radiation.

Additional novelty of this work lies in the establishment of the endoluminal RC model that recapitulates clinical response to therapy. This model builds on our recent murine model of transplantation^18^, but is also the first demonstration that treated and untreated human RC tumors can engraft within the mouse rectum and recapitulate the process of primary disease formation, subsequent invasion, and eventual metastasis. Notably, we observed the development of metastases that may signal tropism of the human RCs from which the tumoroids were derived. While these findings are intriguing, they require prospective validation in a larger cohort. Overall, these data indicate that we have established a working RC platform to better elucidate RC pathogenesis and address questions regarding chemosensitivity and radiation resistance.

In summary, our work addresses an unmet need by establishing a rectal cancer-specific *ex vivo* and *in vivo* biorepository that markedly increases the ability to study rectal cancer. Our model molecularly resembles RC and establishes a relevant framework in which to study the disease. This methodology demonstrates options for eventual drug screening^33^ in a pre-clinical setting and reflects the clinical outcomes of the patients from which they were derived. Our study demonstrates determination of basic response parameters within weeks of derivation and can serve as a robust tool for therapeutic response modeling whereby we can study fundamental research questions relevant to individual RC patients.

## Methods

### Data availability statement

The data that support the findings of this study are available from the corresponding authors upon reasonable request. The data for the RC tumoroids on the cBioPortal will be made publicly available on publication.

### Study Design

We developed a 3D organoid system to culture patient-derived rectal cancer (RC) specimens from treated and untreated patients and dissociated 3D tumoroid and normal adjacent rectal cells (organoids) for long-term use. We developed an in vivo orthotopic, xenograft endoluminal model of rectal cancer with the tumoroids and tested initiation, invasion, tumor growth and chemoresistance in this endorectal endoluminal model.

### Statement of compliance with internal review boards

Animal and human experiments were approved by the following MSKCC Institutional Review Board (IRB) and Institutional Animal Care and Use Committee (IACUC) protocols: #11-083, #16-1071, and 06-07-012. Patients also consented for tissue use and MSK-IMPACT sequencing on the following protocols: #06-107 and #12-245.

### Tissue processing for organoid and tumoroid derivation

For derivation of normal rectal organoids, samples of normal rectal mucosa adjacent to tumor were washed with ice-cold PBS-Abs buffer (PBS with antibiotic-antimycotic, gentamicin [Gemini Bio Product, West Sacramento, CA, USA] and 5 μg/mL metronidazole [SigmaAldrich, St. Louis, MO, USA]). Samples were then chopped into 1 mm pieces in ice-cold PBS-DTT buffer (PBS with 10 mM DTT). The tissue fragments were resuspended in PBS-EDTA (PBS with 8 mM EDTA and 0.5 mM DTT) and were incubated at 4 °C for 1 hr. The suspension was centrifuged at 300×g for 5 min and PBS-EDTA was removed. Tissue fragments were then resuspended vigorously in ice-cold ADF medium (advanced DMEM/F12 plus GlutaMAX, HEPES, and 5% FBS). The tissue fragments were allowed to settle under normal gravity for 1 min, and the supernatant was transferred to a new tube for crypt inspection by microscopy. This step was repeated until no crypts were found in supernatants. The supernatants containing crypts were collected, filtered through a 70 μm Cell Strainer, and then centrifuged at 300×g for 5 min. Isolated crypts were embedded in Matrigel and culture started.

For derivation of tumoroids, fresh rectal cancer samples were processed as reported^21^ and as noted here. Surgically resected rectal tissue or biopsy tissue was washed with ice-cold PBS-Abs buffer and then chopped into 1 mm pieces in ice-cold PBS-DTT buffer. The fragments were digested in digestion medium (advanced DMEM/F12 with 2% FBS, Pen/Strep, 100 U/mL collagenase type XI [SigmaAldrich], and 125 μg/mL dispase type II [Invitrogen, Waltham, MA, USA]) at 37 °C for 40 min and then further digested for 10 min by adding a half-volume of TrypLE Express, and 3 mg of DNase I (SigmaAldrich) per sample. Samples derived from biopsies were then embedded in 800 μL Matrigel and cultured as described below. Samples derived from resected tumors were first filtered; specifically, the tumor cells were then collected by filtering through a 70 μm Cells Strainer and then spinning at 300×g for 5 min. The isolated tumor cells were then embedded in 1-2 mL of Matrigel, depending on pellet size.

### 3D culture conditions for rectal cancer tumoroids

The basal culture medium for healthy tissue-derived organoids and RC tumoroids was modified from the literature (Sato et al. 2011)^21^ as follows: advanced DMEM/F12 was supplemented with antibiotic-antimycotic (Gibco, Waltham, MA, USA), 1×B27, 1×N2, 2 mM GlutaMAX (Invitrogen, Waltham, MA, USA), 10 nM gastrin I, 10 mM HEPES, 1 mM N-acetylcysteine, and 10 mM nicotinamide (SigmaAldrich, St. Louis, MO, USA). The following niche factors were used: 50% Wnt-3A conditioned medium, 20% R-spondin conditioned medium (media collected from HEK293 cell lines expressing recombinant Wnt3a and R-spondin1, kindly provided by Kevin P. O’Rourke)^34^, 100 ng/mL mouse recombinant noggin (Peprotech, Rocky Hill, NJ, USA), 50 ng/mL human recombinant EGF (Peprotech, Rocky Hill, NJ, USA), 500 nM A83-01 (Tocris, Avonmouth, Bristol, UK), and 10 μM SB 202190 (SigmaAldrich, St. Louis, MO, USA). Upon expansion, RC tumoroids were passaged and then cultured in medium without Wnt-3A, R-spondin, and noggin.

For passage of organoids and tumoroids, the Matrigel surrounding the organoids or tumoroids was depolymerized by cell recovery solution (BD Biosciences, San Jose, CA, USA). The released cell clusters were pelleted. For further dissociation into single cells, the pellets were resuspended in TrypLE Express with pipetting 50 times using a p1000 or p200 pipette, and then incubated at 37 °C for 5 min. Cells were then pipetted several more times to form a homogeneous resuspension. Fivefold volume of culture medium was added to the suspension. Pelleted cells were obtained by centrifuging at 300×g for 5 min and supernatant was discarded. The pellet was resuspended with Matrigel and divided into a 24-well suspension plate (50 μL/well). After the Matrigel balls were polymerized, 500 μL of culture medium was added.

To freeze organoids or tumoroids, Matrigel was dissolved with cell recovery solution and then the cell clusters were pelleted. The pellets were resuspended in ice-cold Recovery Cell Culture Freezing Medium (Gibco, Waltham, MA, USA). The pellets also could be dissociated with TrypLE express before Freezing Medium was added. The suspension was frozen down slowly at −80 °C at least overnight followed by storage in liquid nitrogen. The frozen cells were recoverable with standard cell recovery protocol^21^, could be embedded in Matrigel, and culture started as described above. Tumoroids were judged to be successful if they met the following criteria: maintenance in culture for > 4 weeks, viability after five serial passages, and ability to resume growth following multiple freeze/thaw cycles.

### Endoluminal tumor injections

Adapting the dextran sulfate sodium endoluminal protocol^18^ and taking advantage of the Mouse Hospital in the Precision Modeling Center at Memorial Sloan Kettering Cancer Center (MSK) (https://www.mskcc.org/research-programs/precision-disease-modeling), we injected tumoroids into the rectums of NOD scid gamma (NSG) mice. As noted, the experimental protocol was approved by MSK’s Institutional Animal Care and Use Committee (IACUC, IRB #06-07-012). The tumoroids were released from Matrigel by using cell recovery solution and then resuspended in ice-cold PBS with 5% Matrigel to a concentration of 4-6×10^6^ viable cells per mL. Mice were pre-treated with 2.5% dextran sulfate sodium salt (Affymetrix Inc. Cleveland, OH, USA) in drinking water for 5 days and then were allowed to recover for 2 days. With mice under anesthesia, 50 μL of tumoroid suspension was injected slowly via anus into the mice using a p200 pipette. The anuses were then sealed by 5 μL of Tissue Adhesive (Vetbond) for 4 hrs and then the Vetbond was removed. The progression of endoluminal tumor was checked every 4 weeks using small animal endoscopy (Karl Storz Endoscope, El Segundo, CA, USA). Evidence of metastatic disease was assessed by gross examination and microscopic evaluation of resected organs as noted below. The GFP signal was detected by using a 525/45 nm BrightLine^®^ Single-band bandpass filter (IDEX Health & Science, LLC, New York, USA).

### 5-FU treatment of mice with endoluminally implanted tumors

RC-MSK-008 tumoroids were injected into the rectum (see **Fig. 4**) of 15 female (sex matched to tumoroid) mice 6-8 weeks of age. Mice were checked for the progression of tumor using endoscopy at 3-4 week intervals. Then, the 14 mice that were successfully engrafted as evidenced by tumor formation noted on endoscopy were randomized by the envelope method into two groups and intraperitoneally injected with either 5-FU (5 mg/mL in PBS, 50 mg/kg/week), or PBS as control for the duration of the experiment. After 7 weeks of treatment, the tumors were assessed using endoscopy. All of the mice were sacrificed, and rectum, colon, cecum, liver, lung, and brain were isolated and fixed in formalin overnight. Separately, RC-MSK-002 tumoroids were injected into 15 male (sex matched to tumoroid) mice 6-8 weeks of age (see **Fig. 4**). 3 mice did not survive DSS treatment due to severe colitis, and of the remaining 12 mice, 11 successfully engrafted and were treated similarly as with the RC-MSK-008 experiment above.

To quantify tumor size, two independent observers reviewed all endoscopy videos and took screenshots when the endoscope had the tumor in full view. The field of view and tumor were circled and the areas were measured using ImageJ software (NIH). When there were multiple tumors, the sum of the total tumor area was taken. These data are presented as (Area of tumor)/(Area of the field of view) ×100%.

### Ectopic tumor injections

The tumoroids were released from Matrigel by using cell recovery solution and then resuspended in 50% ice-cold PBS and Matrigel (product #356237) solution to a concentration of 1×10^6^ viable cells per mL. Mice under anesthesia were subcutaneously inoculated with 1×10^5^ viable cells in the both hind flanks (100 μL/site). For the RC-MSK-001 experiment, tumors were grown for 6 weeks and for the RC-MSK-002 experiment, tumors were grown for 7 weeks. After this time, the mice were randomized into two groups and intraperitoneally injected with either 5-FU (50 mg/kg), or PBS as control, twice per week for the duration of the experiment. Tumor sizes were measured weekly using a handheld imaging device (Peira TM900).

### Development of the GFP-labeled tumoroid and endoscopic viewing of GFP

The RC-MSK-001-eGFP/luc tumoroids used for bioluminescent tracking were developed by lentivirally introducing GFP and firefly luciferase. Lentiviral particles were generated by transfecting HEK293T cells with the Ubc-eGFP-Luc, psPAX2, and VSV-G^35^ constructs. 7.25 × 10^6^ HEK293T cells were seeded into a 10 cm dish. 7.7 µg of the lentiviral construct, 5.8 µg of psPAX2, and 3.9 µg of VSV-G were transfected using Lipofectamine 2000 according to the manufacturer’s protocol (Invitrogen), grown overnight, and then the medium was replaced with standard DMEM supplemented with 10% FBS, GlutaMAX, and PenStrep. Two days after transfection, the virus media was concentrated using PEG-it Virus Precipitation Solution (SBI System Biosciences), and then the lentiviral particles resuspended in 300 µL PBS. For RC-MSK-001 tumoroid infection, tumoroids from three 50µL Matrigel discs per viral construct were dissociated and resuspended in 10 µL of infection medium (tumoroid culture medium plus 8 µg/mL Polybrene [SigmaAldrich], and 10 µM Y27632 [SigmaAldrich]). The tumoroid cluster suspension and viral suspension were combined in a 48-well culture plate, centrifuged at 600×g at room temperature for 60 minutes, and then incubated for 6 hrs at 37 °C and 5% CO_2_. The contents of each well were homogenized, transferred to a 1.5 mL tube, and centrifuged at 400×g. The supernatant was discarded, the pellet resuspended in 150 µL Matrigel, and the suspension divided into three wells of a 24-well suspension plate. After Matrigel polymerization, 500 µL of culture medium with 10 µM Y27632 was added. Two days after infection, the medium was replaced with tumoroid culture medium. The infected cells were selected for by addition of puromycin (2µg/mL) for 6 days, and then further enriched by fluorescence-activated cell sorting for GFP-positive cells.

### Preparation of tumoroids and tissues for immunohistochemistry and immunofluorescence

After removal of culture medium, the Matrigel disc was washed with PBS, and then fixed with 4% paraformaldehyde for 30 min at room temperature. Then paraformaldehyde was removed, and the fixed sample was stained with eosin and transferred carefully to an embedded cassette. Paraffin embedding and sectioning were processed with standard protocols. Tissues were fixed in formalin and embedded in paraffin. 5 µm sections were used for analysis.

### Immunohistochemistry and immunofluorescence

Hematoxylin and eosin stains were performed on all specimens for initial histopathological evaluation. The immunohistochemistry/immunofluorescence detection was performed at Molecular Cytology Core Facility of Memorial Sloan Kettering Cancer Center using a Discovery XT processor (Ventana Medical Systems), as described in Yarilin et al^36^. Briefly, tissue sections were blocked for 30 minutes, incubated with primary antibody for 4 hrs, and then incubated with biotinylated secondary antibody for 30 minutes. Blocker D and Streptavidin-HRP were used according to the manufacturer instructions (Ventana Medical Systems). Then, for immunohistochemistry, DAB detection kit (Ventana Medical Systems) was used; for immunofluorescence, specimens were incubated with Tyramide-Alexa Fluor 488 (Invitrogen, cat. #T20922) or 568 (Invitrogen, cat. #T20914).

The following antibodies were used: rat anti-mouse EPCAM (eBiosciences, cat. #14-5791), rabbit anti-human EPCAM (Cell Signaling, cat. #2626), rabbit anti-collagen IV (Serotec, cat. #2150-1470), mouse anti-human E-cadherin (BD Transduction, cat. #610181), mouse anti-human CK20 (Dako, cat. #M7019), mouse anti-human CDX2 (BioGenex, cat. #MU392A-UC), rabbit anti-human Muc-2 (Santa Cruz, cat. #15334), and rabbit anti-human Ki67 (abcam, cat. #ab16667). The following secondary antibodies were used: goat anti-rabbit IgG (Vector labs, cat. #PK6101), rabbit anti-rat IgG (Vector, cat. #BA-4000), anti-mouse secondary (Vector Labs, MOM Kit BMK-2202). Immunoglobulins specific for each species were used as negative control.

### Imaging

Slides were scanned using the Pannoramic Flash slide scanner (3DHISTECH) using a 20x 0.8 NA objective (Carl Zeiss). Images were examined and representative areas exported using CaseViewer 2.2 (3DHISTECH).

### Alu quantitative PCR

To quantitatively characterize the metastases noted within the mouse lungs and livers as human cells originating from the rectal xenografts, quantitative polymerase chain reaction (q-PCR) was performed with primers specific for human Alu repeats^37^ and mouse short interspersed nuclear elements (SINE)^38^. This was done on the lung and liver metastases along with a NSG normal adjacent rectal control and a human colon control. The following primers were used: hAlu-101F: 5’-GGTGAAACCCCGTCTCTACT-3’; hAlu-206R: 5’-GGTTCAAGCGATTCTCCTGC-3’; mSINE-F: 5’-AGATGGCTCAGTGGGTAAAGG-3’; mSINE-R: 5’-GTGGAGGTCAGAGGACAAACTT-3’. The reaction was performed in a final volume of 20 uL using FAST SYBR^TM^ Green Master Mix (Applied Biosystems, #4385612) with 0.1 µM of each primer and 20 ng of genomic DNA using an Applied Biosystems 7500 real-time PCR instrument (ABI 7500; Thermo Fisher Scientific). PCR conditions were as follows: 95 °C for 10 min; 45 cycles of 95 °C for 15 s, 56 °C for 30 s, and 72 °C for 30 s. The relative quantity of human Alu sequences was normalized against the relative quantity of mouse SINE sequence for each respective control as ΔCt = Ct^hAlu^ − Ct^mSINE^ and expressed as 2^−ΔCt^. Human and mouse genomic DNA were used as positive and negative controls, respectively. All Alu Ct values were normalized to human DNA Ct value as 100% (human normal adjacent colon TS62T).

### DNA isolation

Tumoroids and normal organoids were released from Matrigel by using cell recovery solution. DNA was extracted using DNeasy Blood and Tissue kit according to the manufacturer’s procedure (Qiagen, Germantown, MD, USA).

### Genomic Analysis

DNA derived from the tumor biopsies and the cultured tumoroids was subjected to exon capture sequencing using the MSK Integrated Mutation Profiling of Actionable Cancer Targets (MSK-IMPACT) platform as previously described^39^. Matched germline DNA samples, extracted from blood and cultured normal organoids, were also sequenced to identify somatic versus germline nucleotide variants. Somatic variants were called using MuTect (v1.1.4) for single nucleotide variants and Somatic Indel Detector (GATK 2.3-9) for indels. Somatic variants were annotated by Annovar for cDNA and amino acid changes and for their presence in dbSNP (v137), the COSMIC database (v68), and 1,000 Genomes minor allele frequencies. Variants were also annotated for their presence and predicted oncogenic status in OncoKB^40^. Each locus where a variant was called was genotyped in matched timepoint samples from the same patient. If a variant was called in one sample and was present in another sample from the same patient in at least 3 reads and at least 1% of reads, it was marked as ‘detectable’ in the other sample. We considered the most frequently mutated genes in the tumoroids and compared their mutation frequency within the tumoroid cohort with the mutation frequency in a set of 291 prospectively analyzed rectal adenocarcinoma samples from MSK. These data can be viewed on the cBioPortal for Cancer Genomics^41,42^. In addition, we used the FACETS algorithm to determine allele specific copy number and cancer cell fraction of each mutation in tumoroids and their respective tumors^43^. Mutations with higher or equal to 85% cancer cell fraction were categorized as clonal. Others were categorized as subclonal.

### Drug treatments

Tumoroids were resuspended in Matrigel and embedded in suspension in a 24-well plate (2-5×10^4^ cells/50 μL Matrigel/well) or 48-well plate (2-3 ×10^3^ cells/20 μL Matrigel/well). The cells were allowed to recover for 2 to 3 days. Medium was replaced with fresh culture medium with varying concentrations of the drugs as follows. For cetuximab treatments, *KRAS*^mutant^ and *KRAS*^wild type^ tumoroids were cultured with cetuximab at 20, 10, 5 and 2.5 μg/mL for 72-hours. For 5-FU treatments, tumoroids were cultured with 5-FU at 500, 100, 50, 10, 5, 1, and 0.5 μM for 6 days, replenishing 5-FU medium on day 3. At the end of treatment in both experiments, the number of viable cells was determined by MTS assay or CellTiter-Glo assay (Promega, Madison, WI, USA) following the kit protocol. IC_50_ values were determined using the Variable Slope Model to calculate a four-parameter dose-response curve using GraphPad Prism 7 (La Jolla, CA, USA).

### CEA assessments

CEA was measured clinically (ng/mL) over the course of treatment from baseline for the patients from which the 6 individual RC tumoroids were derived as noted in **Figure 2b**. The CEA values from baseline and post-treatment were retrieved from the medical record for each patient from which the tumoroids were derived in **Figure 2**. Percent change from baseline was reported as described in the results.

### Irradiation experiments

The tumoroid samples were given to our collaborators (MA, PBP) without supplying any information on the individual patients’ clinical courses. Irradiation was delivered with a Shepherd Mark-I unit (Model 68, SN643, J. L. Shepherd & Associates, San Fernando, CA) operating Cs-137 sources at 1.72 Gy/min (RNK laboratory). Tumoroids were passaged one day prior to the experiment and plated at a density of 100-150 organoids/well in at least 3 wells for each dose. Tumoroids were exposed to single fraction radiation (range 0-20Gy). Six days after irradiation, surviving organoids were manually counted under a brightfield microscope classic dose-response curves were generated. The survival fraction was calculated as the number of surviving tumoroids/number of tumoroids in the un-irradiated sample. Radiation dose survival curves were fitted to the Single Hit Multi Target (SHMT) model^44^ using GraphPad Prism 6. This model allows calculation of D_0_ (the dose required to reduce the fraction of surviving tumoroids to 37% of its previous value) and D_q_ (a threshold dose below which there is no effect) as described^44^.

### Statistics

Statistical analyses reported in **Figure 2a** were calculated using the Kruskal-Wallis test for nonparametric data on each individual 5-FU concentration. The Kruskal-Wallis test statistic (H), degrees of freedom (DF) and P-value for each test are reported as follows and the P-value per concentration is shown in the **Figure 2** legend: <u>0.01 μM</u>: H = 0.8874, DF = 20, P = 0.971; <u>5 μM</u>: H = 8.12, DF = 23, P = 0.150; <u>10 μM</u>: H = 10.3, DF = 23, P = 0.067; <u>50 μM</u>: H = 16.58, DF = 23, P = 0.005; <u>100 μM</u>: H = 14.31, DF = 23, P = 0.014; <u>500 μM</u>: H = 16.36, DF = 23, P = 0.0059). The correlation analysis reported in **Figure 2b** was calculated using Pearson’s correlation. Statistical analysis relating to **Figure 4b** and **4d** were conducted using a 2-tailed student’s t-test with α = 0.05. All error bars for graphical analysis are defined in the respective figure legends.

## Acknowledgments

We thank Brett Carver, M.D. for critical review of the manuscript as it developed. We thank Terry Helms, Medical Illustrator in the Department of Communications at MSK for her expert work in creating illustrations for this work. We thank Dr. Achim Jungbluth for initial discussion on tissue fixation and immunostaining of the murine rectal tissues. We thank Sarat Chandarlapaty, M.D., Ph.D. for discussions regarding the chemoresistance assays and cetuximab resistance assay. We thank Luis Diaz, M.D. for critical comments as this work developed. We thank Nancy Kemeny, M.D. for advice, guidance and encouragement as this project developed. We thank Ning Fan, Sho Fujisawa, Xirong (Frank) Liu, Mesruh Turkekul, and the Molecular Cytology Core for expert assistance and critical feedback in the design and execution of tissue embedding, immunostaining, and processing of tissues in addition to microscopy assistance. We thank the MSK Molecular Core Cytology Facility for critical technical assistance in performing tissue sections, immunohistochemical stains, scans, and analysis (Institutional Core Grant Number P30CA008748). We thank Ronak Shah for initial discussions on set-up of the MSK-IMPACT experiments. We thank Hyun S. Park and Lik Hang Lee, M.D. for assistance with preparation and processing of tumor and tumoroid specimens for IHC and immunofluorescence. We thank Dr. Philip Watson for sharing the Ubc-eGFP-Luc vector.

We also thank members of the Sawyers laboratory for critical review, lively discussion, and helpful comments on this work as it developed. In addition, we thank R. Daniel Beauchamp, M.D. and James R. Goldenring, M.D., Ph.D. for constructive criticism and comments in the development and presentation of this work.

The authors thank Janet Novak, Ph.D. from Memorial Sloan Kettering for editorial assistance.

## Author Contributions

JJS conceived the initial idea behind this work in concert with KG, CW, CLS, and JGA, and edited/wrote the paper with KG/CW/CLS. CW, KG, MA, KPO, and JJS performed the experiments and collected the data. JJS, CW, CLS, BCS, and KG made final edits, figures and completed the paper. KG, KPO, SWL, and LED provided the initial technical expertise to complete this work and assisted in editing the paper, along with PR. WRK and HC provided initial expertise and critical input on the derivation of the RC tumoroids, along with critical input on the final figures and methods. MA did the radiation biology experiments with supervision from RNK, PBP, and JJS. IW, RP, AE, JSS, JS and BCS made significant contributions to obtaining and interpreting the data from **Figures 2 and 3**. MS, YZ, HHW, and RP made significant contributions relative to either obtaining or interpreting the data from Figures 1 and Extended Data Figures 4 and 5. SC, CC and JWH played critical roles in the collection, assessment, analysis and execution of the experiments completed in **Figures 1, 3 and 4**. AB, KM-T, and JS played critical roles in the characterization of the model with histopathologic and immunochemical expertise. MRW, RPD, GMN, JGG, AC, RK, EP, IW, PBP, JGA, LBS, JM, and RY contributed patients, critical clinical information, and critiqued and edited the paper. MB supervised MS, HHW, and YZ, and conceived the MSK-I experiments and data interpretation with JJS. SWL, JGA, and CLS supervised JJS and provided critical input, resources, critique, and oversight of this work.

## Extended Data Figure Legends

**Extended Data Fig. 1 | Rectal cancer tumoroid derivation and patient characteristics. The diagram shows the outcome of attempts to derive tumoroids from** 42 rectal cancer (RC) tumor samples from 25 individual RC patients. 32 RC tumoroids (76%) were successfully derived. For the 10 failed derivations, the points of failure are shown. Demographics from each group are displayed (RAS status [wildtype (WT) or mutant (MUT)], neoadjuvant therapy, metastatic status at derivation, location of tumor where the tumoroid was derived [middle/distal or upper rectum], sex, and age). All patients were mismatch repair proficient (not shown).

**Extended Data Fig. 2 | Preservation of rectal cancer histopathology in tumoroids. a**, Gross resected rectal specimen from which the first RC tumoroid line (RC-MSK-001) was derived and brightfield microscopy of the tumoroid in 3D culture 2 months after processing. Lower panels show hematoxylin and eosin (H&E) staining of the patient tumor (bottom left panel) and the derived tumoroid RC-MSK-001 (bottom right panel) in 3D culture, demonstrating conservation of histopathologic characteristics. Scale bars, 50 μm. **b**, Hoechst and cytoplasmic stains of a representative section of the RC-MSK-001 tumoroid demonstrate the luminal and glandular structure. Scale bars, 20 µm. **c**, Perineal recurrence of the original RC-MSK-001 tumor and the tumoroid (RC-MSK-001PR) derived from it is shown. H&E of recurrent tumor and the derived tumoroid are shown. Scale bar, 50 μm. **d**, H&E comparison of seven tumoroid cell lines as noted with the corresponding primary tumor from which they were derived. Scale bars, 50 μm.

**Extended Data Fig. 3 | Conservation of enterocyte markers.** RC-MSK-001, RC-MSK-002 and RC-MSK-008 tumoroids are compared to the respective primary tumors for Alcian blue, CK20, CDX2, MUC-2, and E-cadherin staining. For immunofluorescent staining: E-cadherin (green), DAPI (blue). See **Figure 1a** and **Figure 1b** for another example of RC-MSK-001 H&E, Alcian blue, CK20, and CDX2 comparisons. Scale bars, 50 μm.

**Extended Data Fig. 4 | The mutational fingerprint in derived RC tumoroids. a**, The mutational fingerprint of 15 RC tumoroids for the 50 most common alterations as determined by MSK-IMPACT. The frequency of alteration is noted along with the type of genetic alteration relative to truncating mutation, inframe mutation, missense mutation, or splice site alterations (as noted by the color code). **b**, Example of an individual tumoroid (RC-MSK-003) with complete conservation of mutations between the tumoroid and the primary tumor from which it was derived. **c**, Example of another tumoroid (RC-MSK-004) with conservation of driver mutations and the addition of two secondary mutations that were noted in the tumoroid in culture only.

**Extended Data Fig. 5 | Rectal cancer tumoroids maintain representation of mutations found in the patient tumor. a**, All mutations called in the MSK-IMPACT sequencing of tumoroids and primary tumors are shown. The numbers of mutations are displayed with regard to each gene (by column) and each tumoroid and tumor pair (by row). Mutations are colored by concordance status. **b**, Percentage of concordance between tumoroid and tumor among mutations predicted to be oncogenic overall and by each patient. The mutations represented are those annotated by OncoKB^40^ as oncogenic or likely oncogenic in each tumoroid and tumor pair.

**Extended Data Fig. 6 | Resistance to a targeted anti-epidermal growth factor receptor therapy, cetuximab, in KRAS^mutant^ compared with KRAS^wild type^ tumoroids**. **a**, Resistance to cetuximab is demonstrated in KRAS^mutant^ RC tumoroids compared with a KRAS^wild type^ tumoroid. Dose range was used as shown and percentage of live cells is displayed for each tumoroid. **b**, Independent assay comparing different KRAS^mutant^ and KRAS^wild type^ rectal cancer tumoroids. Similar methodology was used as in panel a. Data are in quadruplicate for panels a and b (error bars = standard error of the mean).

**Extended Data Fig. 7 | Demonstration of endorectally implanted human rectal cancer into a mouse rectum via endoscopic biopsy (see Supplementary Movie 1)**. Serial sections for a mouse implanted with RC-MSK-001 tumoroid stained by IF for human EpCAM (green); collagen IV (red); merged with DAPI (blue). Colon from an unimplanted NSG mouse was used as control. Scale bars: leftmost and middle images, 200 µm; rightmost images, 50 µm.

**Extended Data Fig. 8 | Endoluminal tumors can be assessed by serial endoscopy, immunofluorescence and GFP. a**, 12-week endoscopy of a mouse transplanted with RC-MSK-001 rectal tumoroids. **b**, 12-week endoscopy of a mouse transplanted with the RC-MSK-002 rectal tumoroids. **c**, Representative human (h) and mouse (m) EpCAM staining for an NSG mouse engrafted with RC-MSK-001 tumoroids. Distinct staining was noted for hEpCAM (green) and mEpCAM (red). Scale bars: low magnification, 500 μm; high magnification inset, 50 μm. **d**, Representative hEpCAM and mEpCAM staining for an RC-MSK-002 engrafted NSG mouse. As in panel c, distinct staining was noted for human and mouse EpCAM. Scale bars: low magnification, 500 μm; high magnification inset, 50 μm. **e**, RC-MSK-001 tumoroids labeled with GFP and viewed by brightfield (left panel) and intravital GFP imaging (middle panel). Scale bar, 100 µm. Endoscopic view of these tumoroids transplanted into an NSG mouse is shown (right panel). **f**, Invasive rectal tumor after RC-MSK-001 tumoroid implantation. The micrographs show H&E, immunohistochemistry (IHC) for GFP, and IF stains on serial sections for hEpCAM (green), mEpCAM (red), and human Ki67 (green), each merged with DAPI (blue). Unless otherwise noted, scale bar = 100 μm.

**Extended Data Fig. 9 | The endoluminal rectal cancer model recapitulates invasive cancer. a**, Shown is an independent experiment (from Figure 3c) of a male NSG mouse sacrificed at 22 weeks post-transplantation. Gross implanted rectal tumor, H&E and immunofluorescence are shown. IF serial sections were stained with hEpCAM (green), mEpCAM (red), and DAPI (blue) and show engraftment and invasion of human tumoroids. Scale bars: H&E, 500 μm; IF, 100 μm. **b-d**, An endorectal tumor (RC-MSK-001) is shown 16 weeks after endoluminal transplantation. H&E demonstrates invasion at the junction between the columnar and squamous epithelium of the anorectal junction. Staining is as follows: **b/c**, H&E; **d**, DAPI (blue) + hEpCAM (green) + collagen IV (red); Scale bars are as follows: **b**, 1,000 μm; **c-d**: low magnification, 400 μm; high magnification insets, 100 μm.

**Extended Data Fig. 10 | The rectal cancer endoluminal transplantation assay recapitulates metastases to the liver in a manner similar to the patient. a**, Liver metastasis in an independent experiment similar to that shown in **Figure 3d** is shown after an endoluminal rectal transplantation experiment in a male NSG mouse that was sacrificed at 36 weeks post-transplantation. Gross rectal tumor is shown along with gross liver tumor. Liver tumor is marked by poorly differentiated histology. Scale bars for H&E images, from low to high magnification, are as follows: 1,000 μm, 500 μm and 100 μm**. b**, Axial and coronal CT images of liver metastases in the corresponding patient (arrowheads). **c,** Human-specific Alu qPCR demonstrates that the metastases noted in the mice (**Figure 3d** and current **Extended Data Figure 10**) arose from the implanted human rectal tumoroids in the mouse rectums. Not shown: no signal when using mouse colon indicating the specificity of the assay for human tissue. All data shown in triplicate, error bars = standard deviation (SD).

**Extended Data Fig. 11 | Tumorigenicity in ectopic models and responses to 5-FU therapy *in vivo*. a**, Growth of tumors established from flank injection of RC-MSK-001 tumoroid. The graph displays relative tumor volume over time in the vehicle (n = 4) and 5-FU (n = 6) treated groups. Error bars = SD. **b**, RC-MSK-002 tumoroid flank model (similar as in panel a) for vehicle (n = 2) and 5-FU (n = 3) treated groups. Error bars = SD where shown (if not shown then they were not calculated due to small sample size).

**Extended Data Fig. 12 | Histopathologic conservation of glandular architecture in the endoluminally implanted RC tumoroids**. H&E images are shown for RC-MSK-008 and RC-MSK-002 tumoroids in the endoluminal transplantation model (left) along with their corresponding patient primary tumors (right). The H&E photomicrographs demonstrate histopathologic conservation of glandular features as noted in the human adenocarcinomas from which they were derived. Scale bar, 50 μm.

