## Extended Data Figures for "A rectal cancer model establishes a platform to study individual responses to chemoradiation"

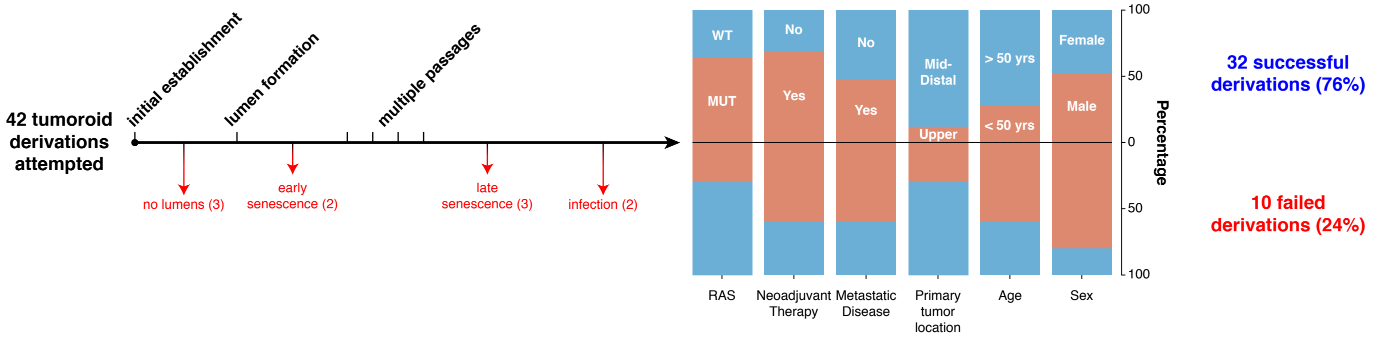


**Extended Data Fig. 1 | Rectal cancer tumoroid derivation and patient characteristics.** The diagram shows the outcome of attempts to derive tumoroids from 42 rectal cancer (RC) tumor samples from 25 individual RC patients. 32 RC tumoroids (76%) were successfully derived. For the 10 failed derivations, the points of failure are shown. Demographics from each group are displayed (RAS status [wildtype (WT) or mutant (MUT)], neoadjuvant therapy, metastatic status at derivation, location of tumor where the tumoroid was derived [middle/distal or upper rectum], sex, and age). All patients were mismatch repair proficient (not shown).

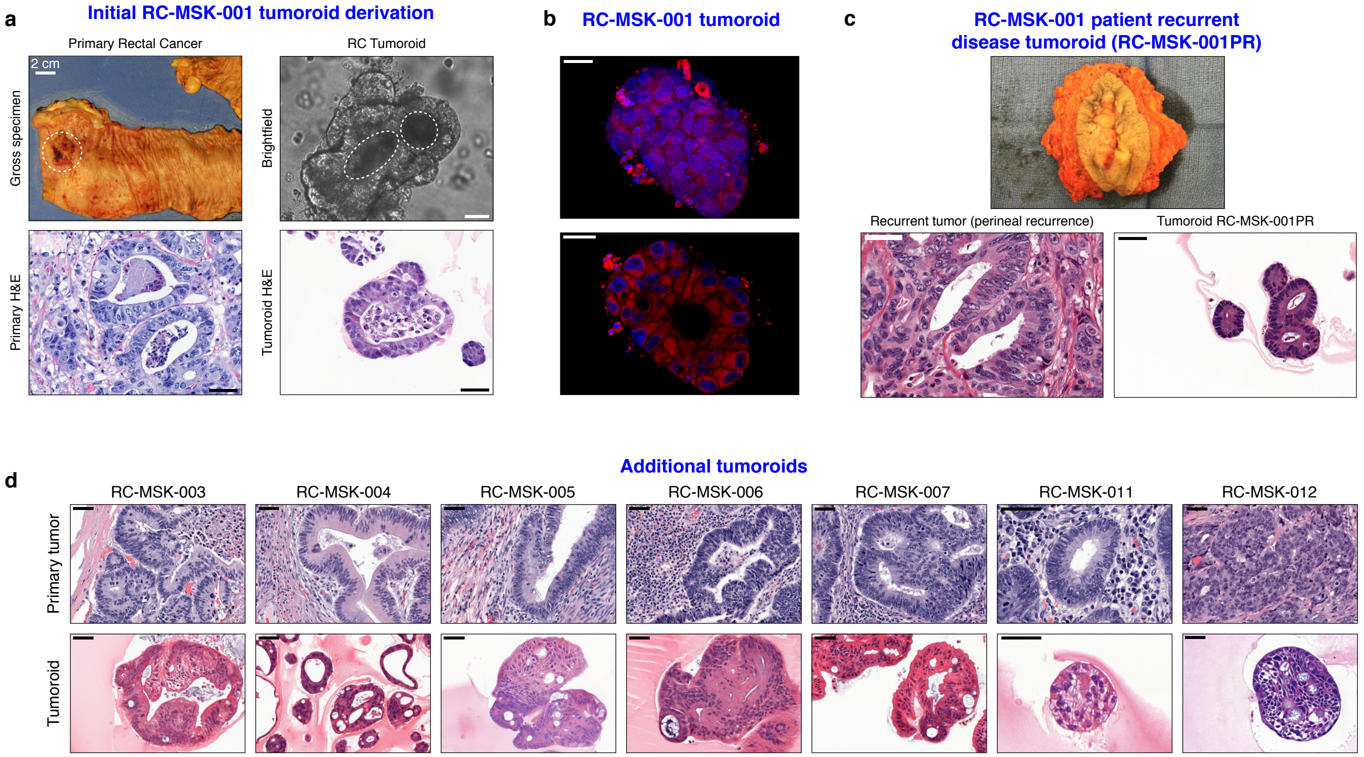


**Extended Data Fig. 2 | Preservation of rectal cancer histopathology in tumoroids.** **a**, Gross resected rectal specimen from which the first RC tumoroid (RC-MSK-001) was derived and brightfield microscopy of the tumoroid in 3D culture 2 months after processing. Lower panels show hematoxylin and eosin (H&E) staining of the patient tumor (bottom left panel) and the derived tumoroid RC-MSK-001 (bottom right panel) in 3D culture, demonstrating conservation of histopathologic characteristics. Scale bars, 50 µm. **b**, Hoechst and cytoplasmic stains of a representative section of the RC-MSK-001 tumoroid demonstrate the luminal and glandular structure. Scale bars, 20 µm. **c**, Perineal recurrence of the original RC-MSK-001 tumor and the tumoroid (RC-MSK-001PR) derived from it is shown. H&E of recurrent tumor and the derived tumoroid are shown. Scale bar, 50 µm. **d**, H&E comparison of seven tumoroid cell lines as noted with the corresponding primary tumor from which they were derived. Scale bars, 50 µm.

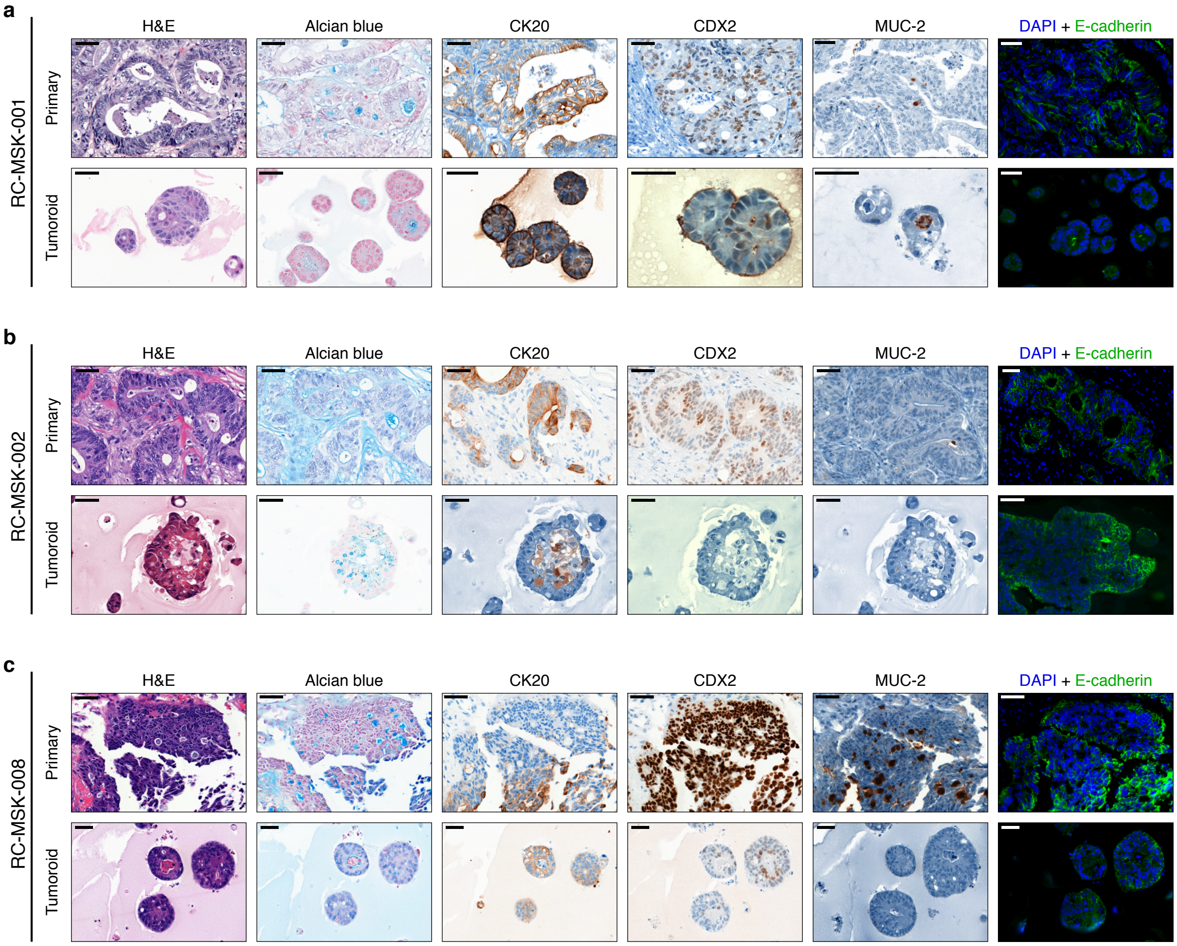


**Extended Data Fig. 3 | Conservation of enterocyte markers.** RC-MSK-001, RC-MSK-002, and RC-MSK-008 tumoroids are compared to the respective primary tumors for Alcian blue, CK20, CDX2, MUC-2, and E-cadherin staining. For immunofluorescent staining: E-cadherin (green), DAPI (blue). See **Figure 1a** and **Figure 1b** for another example of RC-MSK-001 H&E, Alcian blue, CK20, and CDX2 comparisons. Scale bars, 50 µm.

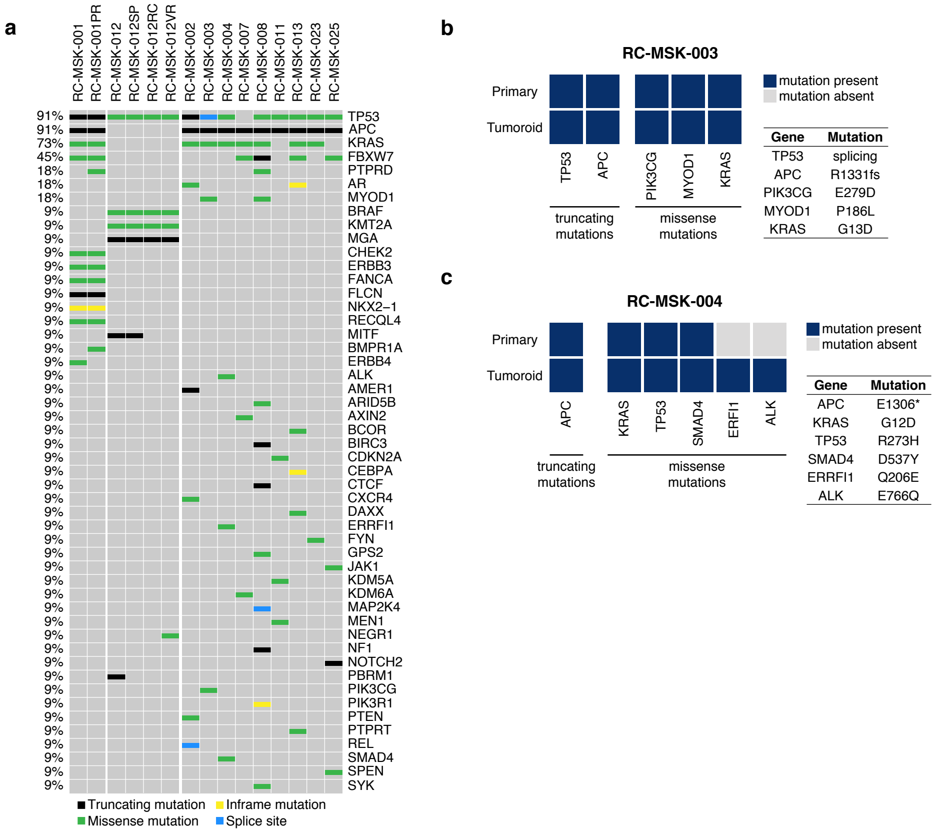


**Extended Data Fig. 4 | The mutational fingerprint in derived RC tumoroids.** **a**, The mutational fingerprint of 15 RC tumoroids for the 50 most common alterations as determined by MSK-IMPACT. The frequency of alteration is noted along with the type of genetic alteration relative to truncating mutation, inframe mutation, missense mutation, or splice site alterations (as noted by the color code). **b**, Example of an individual tumoroid (RC-MSK-003) with complete conservation of mutations between the tumoroid and the primary tumor from which it was derived. **c**, Example of another tumoroid (RC-MSK-004) with conservation of driver mutations and the addition of two secondary mutations that were noted in the tumoroid in culture only.

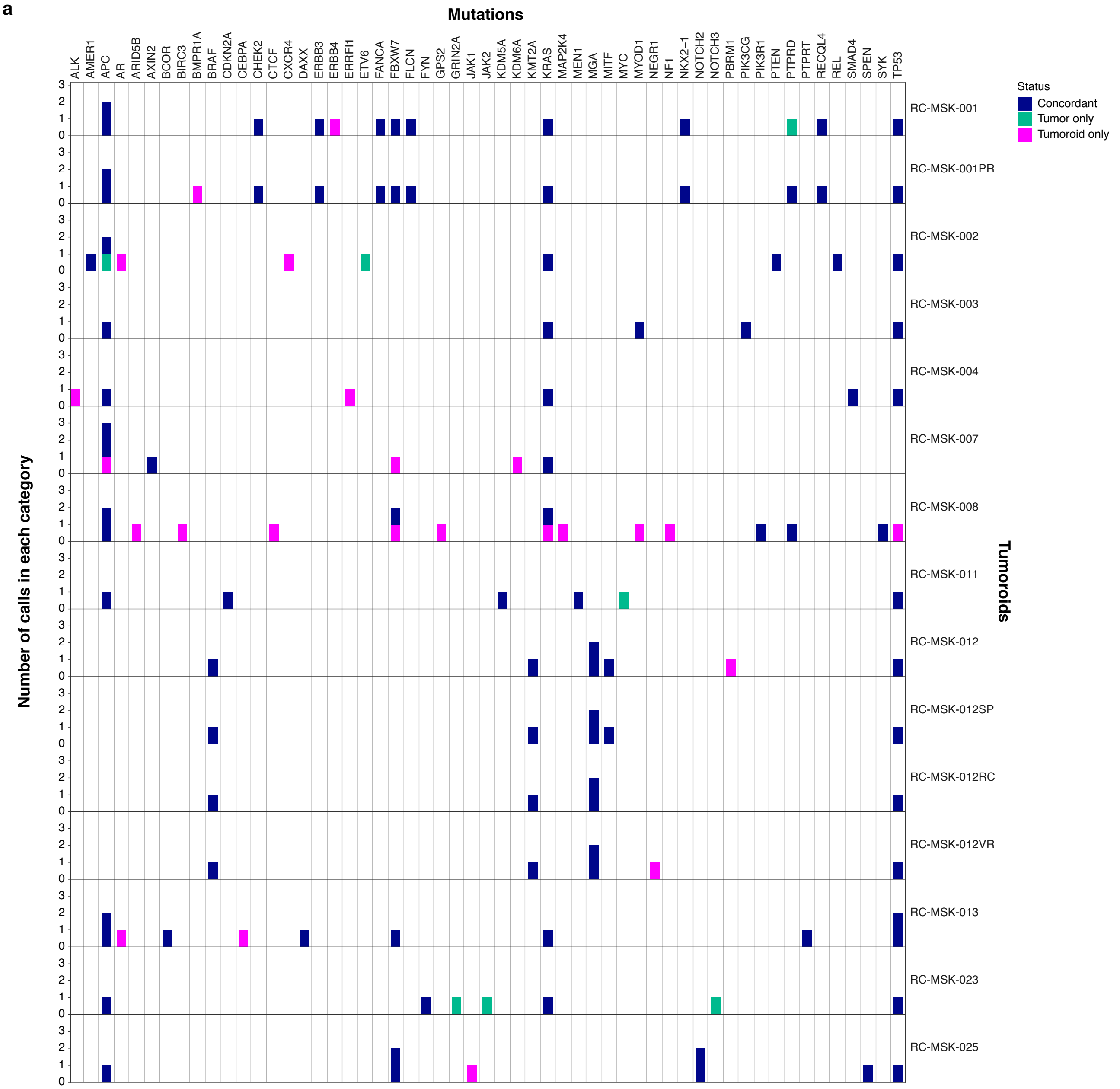


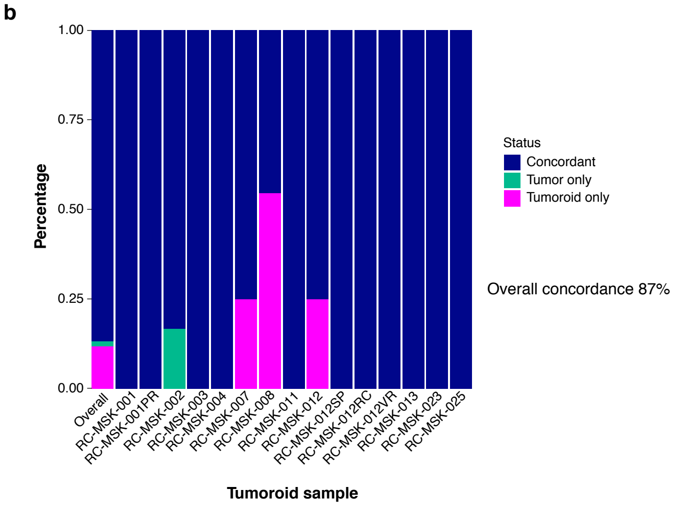


**Extended Data Fig. 5 | Rectal cancer tumoroids maintain representation of mutations found in the patient tumor. a**, All mutations called in the MSK-IMPACT sequencing of tumoroids and primary tumors are shown. The numbers of mutations are displayed with regard to each gene (by column) and each tumoroid and tumor pair (by row). Mutations are colored by concordance status. **b**, Percentage of concordance between tumoroid and tumor among mutations predicted to be oncogenic overall and by each patient. The mutations represented are those annotated by OncoKB[40] as oncogenic or likely oncogenic in each tumoroid and tumor pair.

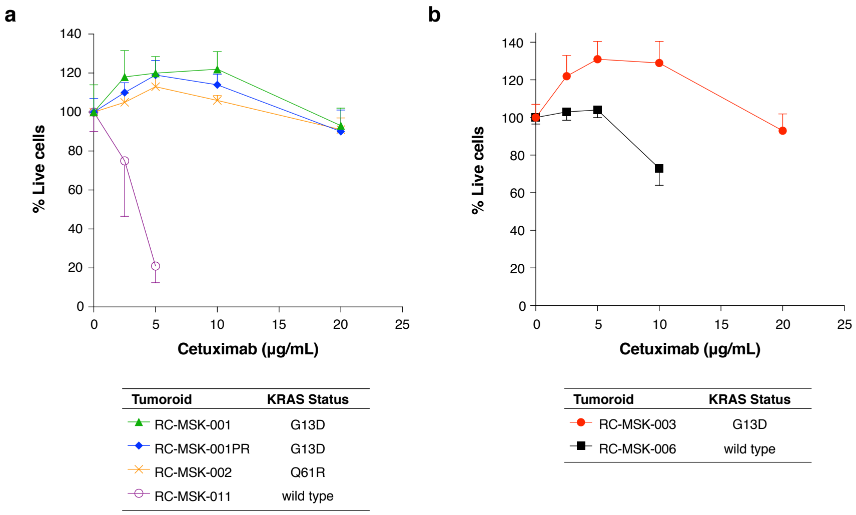


**Extended Data Fig. 6 | Resistance to a targeted anti-epidermal growth factor receptor therapy, cetuximab, in KRAS$^{mutant}$ compared with KRAS$^{wild\ type}$ tumoroids.** **a**, Resistance to cetuximab is demonstrated in KRAS$^{mutant}$ RC tumoroids compared with a KRAS$^{wild\ type}$ tumoroid. Dose range was used as shown and percentage of live cells is displayed for each tumoroid. **b**, Independent assay comparing different KRAS$^{mutant}$ and KRAS$^{wild\ type}$ rectal cancer tumoroids. Similar methodology was used as in panel a. Data are in quadruplicate for panels a and b (error bars = standard error of the mean).

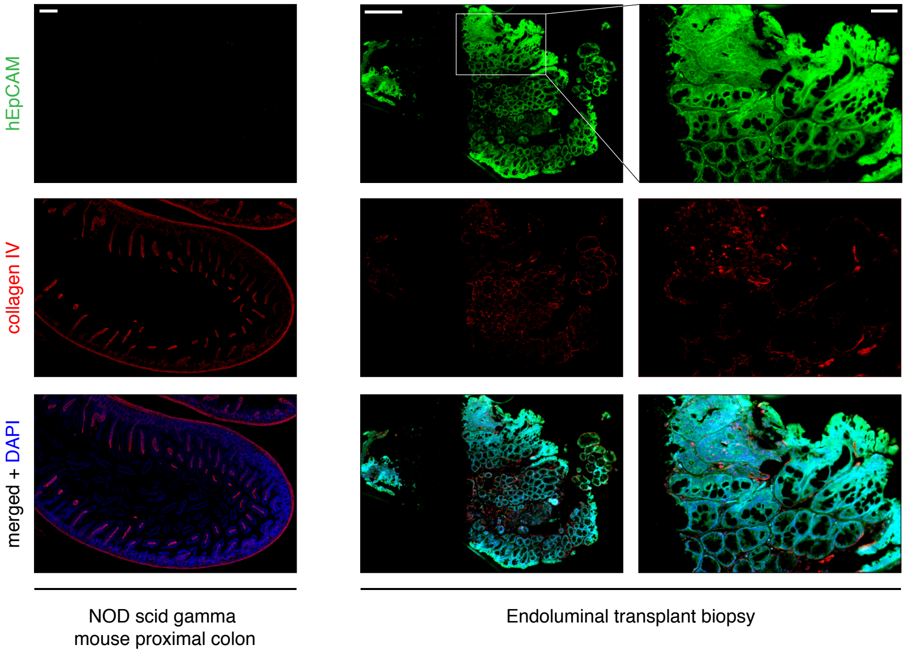


**Extended Data Fig. 7 | Demonstration of endorectally implanted human rectal cancer into a mouse rectum via endoscopic biopsy (see Supplementary Movie 1).** Serial sections for a mouse implanted with RC-MSK-001 tumoroid stained by IF for human EpCAM (green); collagen IV (red); merged with DAPI (blue). Colon from an unimplanted NSG mouse was used as control. Scale bars: leftmost and middle images, 200 µm; rightmost images, 50 µm.

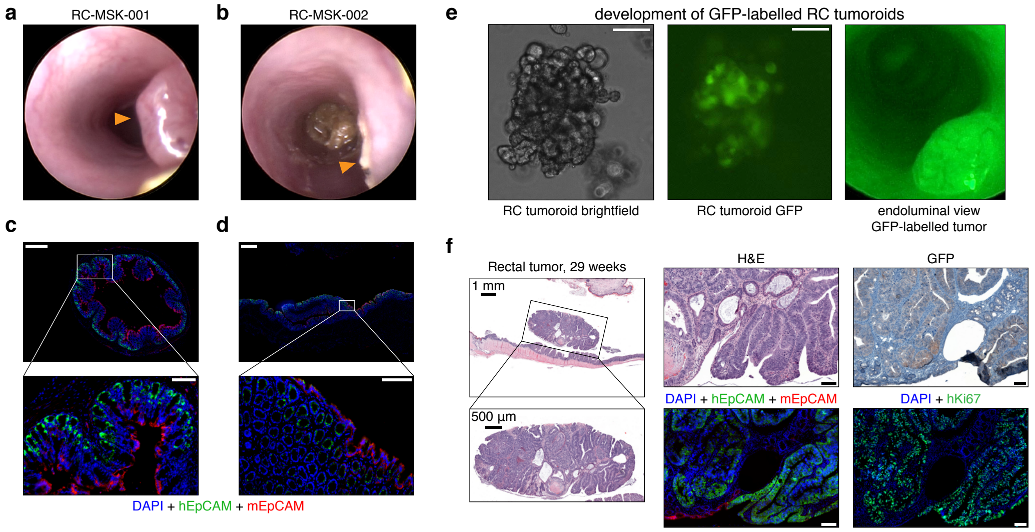


**Extended Data Fig. 8 | Endoluminal tumors can be assessed by serial endoscopy, immunofluorescence and GFP. a**, 12-week endoscopy of a mouse transplanted with RC-MSK-001 rectal tumoroids. **b**, 12-week endoscopy of a mouse transplanted with the RC-MSK-002 rectal tumoroids. **c**, Representative human (h) and mouse (m) EpCAM staining for an NSG mouse engrafted with RC-MSK-001 tumoroids. Distinct staining was noted for hEpCAM (green) and mEpCAM (red). Scale bars: low magnification, 500 µm; high magnification inset, 50 µm. **d**, Representative hEpCAM and mEpCAM staining for an RC-MSK-002 engrafted NSG mouse. As in panel c, distinct staining was noted for human and mouse EpCAM. Scale bars: low magnification, 500 µm; high magnification inset, 50 µm. and viewed by brightfield (left panel) and intravital GFP imaging (middle panel). Scale bar, 100 µm. **e**, RC-MSK-001 tumoroids labeled with GFP. Endoscopic view of these tumoroids transplanted into an NSG mouse is shown (right panel). **f**, Invasive rectal tumor after RC-MSK-001 tumoroid implantation. The micrographs show H&E, immunohistochemistry (IHC) for GFP, and IF stains on serial sections for hEpCAM (green), mEpCAM (red), and human Ki67 (green), each merged with DAPI (blue). Unless otherwise noted, scale bar = 100 µm.

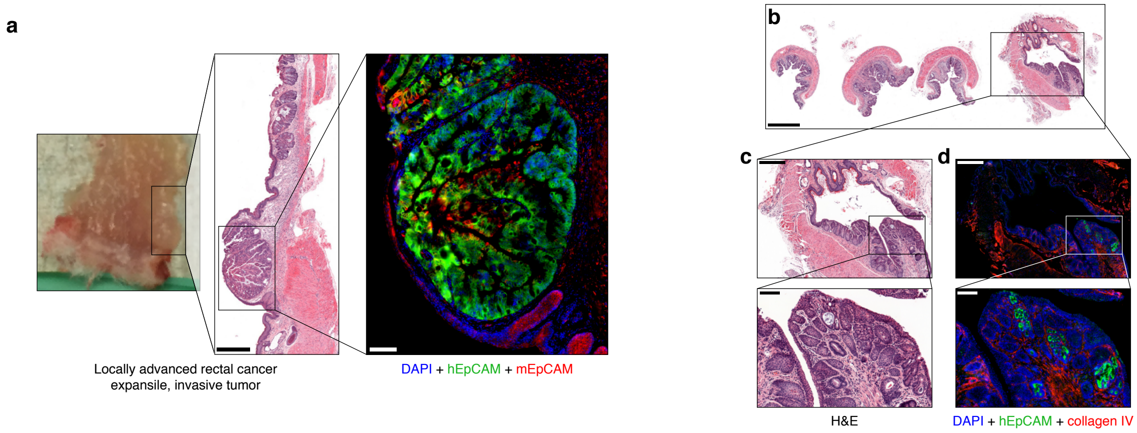


**Extended Data Fig. 9 | The endoluminal rectal cancer model recapitulates invasive cancer. a**, Shown is an independent experiment (from **Figure 3c**) of a male NSG mouse sacrificed at 22 weeks post-transplantation. Gross implanted rectal tumor, H&E and immunofluorescence are shown. IF serial sections were stained with hEpCAM (green), mEpCAM (red), and DAPI (blue) and show engraftment and invasion of human tumoroids. Scale bars: H&E, 500 µm; IF, 100 µm. **b-d**, An endorectal tumor (RC-MSK-001) is shown 16 weeks after endoluminal transplantation. H&E demonstrates invasion at the junction between the columnar and squamous epithelium of the anorectal junction. Staining is as follows: b/c, H&E; d, DAPI (blue) + hEpCAM (green) + collagen IV (red); Scale bars are as follows: b, 1,000 µm; c-d: low magnification, 400 µm; high magnification insets, 100 µm.

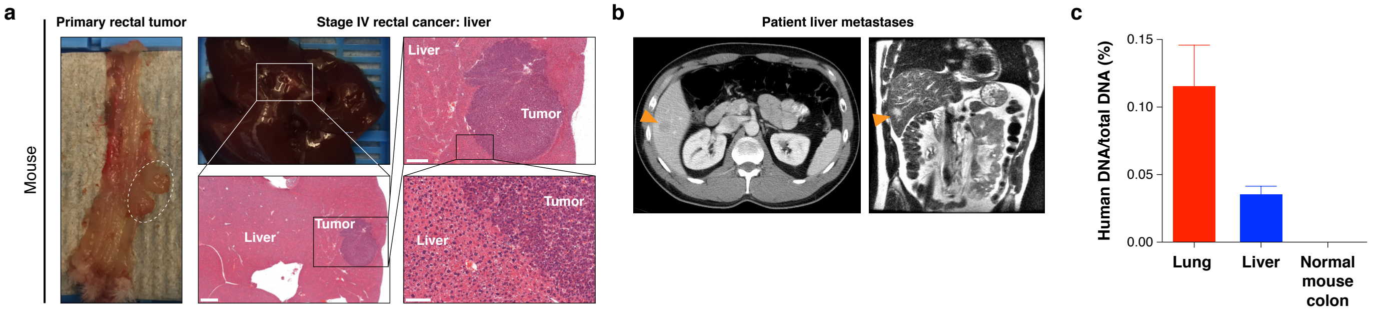


**Extended Data Fig. 10 | The rectal cancer endoluminal transplantation assay recapitulates metastases to the liver in a manner similar to the patient. a**, Liver metastasis in an independent experiment similar to that shown in Figure 3d is shown after an endoluminal rectal transplantation experiment in a male NSG mouse that was sacrificed at 36 weeks post-transplantation. Gross rectal tumor is shown along with gross liver tumor. Liver tumor is marked by poorly differentiated histology. Scale bars for H&E images, from low to high magnification, are as follows: 1,000 µm, 500 µm and 100 µm. **b**, Axial and coronal CT images of liver metastases in the corresponding patient (arrowheads). **c**, Human-specific Alu qPCR demonstrates that the metastases noted in the mice (Figure 3d and current Extended Data Figure 10) arose from the implanted human rectal tumoroids in the mouse rectums. Not shown: no signal when using mouse colon indicating the specificity of the assay for human tissue. All data shown in triplicate, error bars = standard deviation (SD).

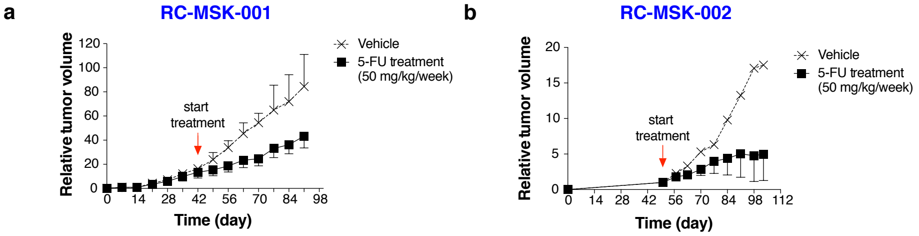


**Extended Data Fig. 11 | Tumorigenicity in ectopic models and responses to 5-FU therapy in vivo. a**, Growth of tumors established from flank injection of RC-MSK-001 tumoroid. The graph displays relative tumor volume over time in the vehicle (n = 4) and 5-FU (n = 6) treated groups. Error bars = SD. **b**, RC-MSK-002 tumoroid flank model (similar as in panel a) for vehicle (n = 2) and 5-FU (n = 3) treated groups. Error bars = SD where shown (if not shown then they were not calculated due to small sample size).

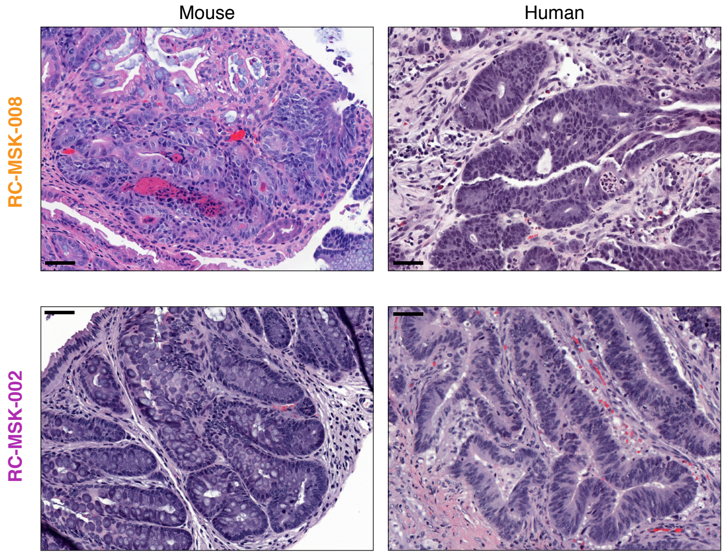


**Extended Data Fig. 12 | Histopathologic conservation of glandular architecture in the endoluminally implanted RC tumoroids.** H&E images are shown for RC-MSK-008 and RC-MSK-002 tumoroids in the endoluminal transplantation model (left) along with their corresponding patient primary tumors (right). The H&E photomicrographs demonstrate histopathologic conservation of glandular features as noted in the human adenocarcinomas from which they were derived. Scale bar, 50 µm.
