## Supplementary Table for "A rectal cancer model establishes a platform to study individual responses to chemoradiation"

**Supplementary Table 1 | Rectal cancer patient and rectal cancer tumoroid characteristics.** All patients had intact mismatch repair.

| Name | Tissue site of derivation | Tissue studies completed | Tumoroid studies completed | Normal organoid isolated | Age at derivation | Sex | RAS status | Neoadjuvant therapy | Treatment response in resected primary | Type of adjuvant therapy | Metastatic disease present at derivation |
|---|---|---|---|---|---|---|---|---|---|---|---|
| RC-MSK-001 | Distal rectum | IHC, IF, MSK-I | IHC, IF, MSK-I, *ex vivo*, *in vivo* | No | 56 | F | KRAS mutant (G13D) | FOLFOX, FOLFIRI, SCRT and FUDR[†] | Incomplete | none | Yes (lung, liver, brain) |
| RC-MSK-001PR | Very distal rectum (recurrence) | IHC, IF, MSK-I | IHC, MSK-I, *ex vivo* | No | 57 | F | KRAS mutant (G13D) | holiday between rectal resection and recurrence[†] | Incomplete | FOLFIRI + Bev | Yes (lung, liver, brain) |
| RC-MSK-002 | Upper rectum/distal sigmoid | IHC, IF, MSK-I | IHC, IF, MSK-I, *ex vivo*, *in vivo* | No | 42 | M | KRAS mutant (Q61R) | none | n/a | FOLFOX | Yes (liver/lung) |
| RC-MSK-003 | Distal rectum | IHC, MSK-I | IHC, MSK-I, *ex vivo* | No | 71 | M | KRAS mutant (G13D) | FOLFOX and SCRT[†] | Incomplete | 5-FU/LV | Yes (liver/perineum) |
| RC-MSK-003PR | Distal recurrence | IHC | IHC | Yes | 72 | M | Not tested | FOLFOX then SCRT[†] | Regrowth after CAA | 5-FU, now FOLFIRI | Yes (lung/liver) |
| RC-MSK-004 | Mid-distal rectum | IHC, MSK-I | IHC, IF, MSK-I, *ex vivo*, *in vivo* | No | 43 | M | KRAS mutant (G12D) | FOLFOX[†] | Incomplete | FOLFIRI, FUDR, and now CapeOx | Yes (liver) |
| RC-MSK-005 | Mid-to-distal rectum (obstructing) | IHC | IHC | No | 52 | F | KRAS mutant (G12A) | FLOX, FOLFOX[†] | Incomplete | None | Yes (lung) |
| RC-MSK-006* | Mid rectum | IHC | IHC | No | 60 | M | Wild type | FOLFOX, CRT[†] | Incomplete | n/a | No |
| RC-MSK-007 | Mid rectum | IHC, MSK-I | IHC, MSK-I | Yes | 66 | M | KRAS mutant (A146T) | FOLFOX, CRT[†] | Incomplete | n/a | No |
| RC-MSK-008 | Distal rectum | IHC, MSK-I | IHC, MSK-I, *ex vivo*, *in vivo* | No | 70 | F | KRAS mutant (G12S) | FOLFOX planned | n/a | n/a | Yes (lung) |
| RC-MSK-009 | Upper rectum | IHC, MSK-I | IHC, MSK-I, *ex vivo* | Yes | 44 | M | Wild type | FOLFOX only (Prospect trial)[†] | Incomplete | 5-FU/LV x 6 | No |
| RC-MSK-010 | Distal rectum | IHC | Brightfield | Yes | 39 | F | Wild type | FOLFOX + CRT[†] | Incomplete | FOLFIRI, Lonsurf, EGFR- | Yes (peritoneum/vagina) |
| RC-MSK-011 | Mid rectum | IHC | IHC | Yes | 62 | M | KRAS mutant (G12D) | FOLFOX and CRT[†] | Incomplete | n/a | No |
| RC-MSK-012 | Mid-to-distal rectum | IHC, MSK-I | IHC, MSK-I, *ex vivo* | Yes | 45 | F | Wild type | FOLFOX, CRT then FOLFIRI[†] | Incomplete | FOLFIRI | Yes (spleen, pelvis) |
| RC-MSK-012SP | Splenic met from primary | IHC, MSK-I | IHC, MSK-I, *ex vivo* | No | 45 | F | Wild type | FOLFOX, CRT then FOLFIRI[†] | n/a | n/a | n/a |
| RC-MSK-012VR | Pelvic recurrence | IHC | IHC, MSK-I, *ex vivo* | No | 45 | F | Wild type | FOLFOX, CRT then FOLFIRI[†] | n/a | FOLFIRI/Bev | Yes |
| RC-MSK-012RC | Pelvic recurrence | IHC | IHC, MSK-I, *ex vivo* | No | 45 | F | Wild type | FOLFOX, CRT then FOLFIRI[†] | n/a | FOLFIRI/Bev | Yes |
| RC-MSK-013 | Distal rectum | IHC, MSK-I | IHC, MSK-I, *ex vivo* | Yes | 84 | F | KRAS mutant (G13C) | CRT then 5-FU planned | POD | n/a | Local POD |
| RC-MSK-014* | Mid-rectal tumor | IHC | IHC | Yes | 70 | F | Not tested | FOLFOX then CRT[†] | cCR then local regrowth (20%) | None | No |
| RC-MSK-015 | Mid-upper | IHC | IHC | Yes | 47 | F | KRAS mutant (Q61R) | CapeOx[†] | cCR then local regrowth (50%) | None | No |
| RC-MSK-016 | Mid to distal | IHC | IHC | No | 86 | F | NRAS | FOLFOX planned | Incomplete | None | No |
| RC-MSK-016PC | Mid to distal | IHC | IHC | No | 86 | F | NRAS | post-FOLFOX | n/a | n/a | No |
| RC-MSK-016RES | Distal rectum | IHC | IHC | Yes | 86 | F | NRAS | Post-FOLFOX and CRT | Incomplete | n/a | No |
| RC-MSK-017 | Upper rectal primary | IHC | IHC | Yes | 81 | F | Not tested | none | n/a | None | Yes (liver, peritoneum) |
| RC-MSK-018 | Mid-rectal tumor | IHC | IHC | Yes | 59 | M | KRAS mutant (G12V) | FOLFOX[†] | n/a (bx) | n/a | Yes (liver) |
| RC-MSK-019 | Mid-rectal tumor | IHC | IHC | Yes | 88 | M | KRAS mutant (K147E) | Capecitabine then CRT planned | incomplete | n/a | No |
| RC-MSK-021 | Mid-rectal tumor | IHC | Brightfield | Yes | 62 | M | Not tested | Capecitabine then SCRT[†] | pCR | None | No |
| RC-MSK-022* | Mid-to-distal Rectum | IHC | IHC, *ex vivo* | Yes | 56 | M | Wild type | FOLFOX then CRT planned | TBD | n/a | No |
| RC-MSK-023 | LAR recurrence (distal) | IHC, MSK-I | IHC, MSK-I, *ex vivo* | Yes | 45 | M | KRAS mutant (Q61H) | FOLFOX, then CRT then combo resection[†] | incomplete | FOLFIRI | Yes (liver and lung) |
| RC-MSK-024 | Mid-rectum | IHC | IHC | Yes | 52 | F | Wild type | FOLFOX then CRT planned | TBD | n/a | No |
| RC-MSK-025* | Mid-rectum | IHC, MSK-I | IHC, *ex vivo* | Yes | 51 | M | Wild type | CRT then FOLFOX planned | nCR | n/a | No |
| RC-MSK-026 | Mid-rectum | IHC | IHC | Yes | 65 | F | Wild type | FOLFOX[†] | Incomplete | FUDR | Yes |

* = tumoroid derived while on trial (RC-MSK-006 on PROSPECT; RC-MSK-014, RC-MSK-022, RC-MSK-025 on OPRA). † = tumoroid derived while on neoadjuvant therapy. - = not yet completed. Tumoroid nomenclature: PR = perineal recurrence; SP = splenic metastasis; VR = pelvic recurrence vagina; RC = pelvic recurrence rectal; PC = post-5-FU based chemotherapy; RES = resection; PM = peritoneal metastasis. Other abbreviations: IHC = Immunohistochemistry; IF = immunofluorescence; MSK-I = MSK Integrated Mutation Profiling of Actionable Cancer Targets, MSK-IMPACT; EA = Endoluminal assay; POD = progression of disease; pCR = pathologic complete response; cCR = clinical complete response; nCR = near complete response; CAA = coloanal anastomosis; SCRT = short-course radiation therapy; CRT = long-course chemoradiation; 5-FU = 5-fluorouracil; LV = leucovorin; FLOX = 5-FU + oxaliplatin; FOLFOX = 5-FU + leucovorin + oxaliplatin; FOLFIRI = 5-FU + leucovorin + irinotecan; CapeOx = capecitabine + oxaliplatin; FUDR = floxuridine; Lonsurf = trifluridine and tipiracil; Bev = bevacizumab.
